# Exosome-transmitted miR-769-5p confers cisplatin resistance and tumorigenesis in gastric cancer by targeting CASP9 and promoting the ubiquitination degradation of p53

**DOI:** 10.1101/2021.09.19.461013

**Authors:** Xinming Jing, Mengyan Xie, Kun Ding, Tingting Xu, Yuan Fang, Pei Ma, Yongqian Shu

## Abstract

Cisplatin resistance is the main cause of poor clinical prognosis in patients with gastric cancer (GC). Yet, the exact mechanism of cisplatin resistance remains unclear. Recent studies have suggested that exocrine miRNAs found in the tumor microenvironment participates in tumor metastasis and drug resistance. In this study, we discovered that cisplatin-resistant GC cells communicate with the tumor microenvironment by secreting microvesicles. The biologically active miR-769-5p can be integrated into exosomes and delivered to sensitive cells, thereby spreading cisplatin resistance. Mi769-5p was upregulated in GC tissues and enriched in the serum exosomes of cisplatin-resistant patients. Mechanistically, miR-769-5p promotes cisplatin resistance by targeting CASP9 so as to inhibit the downstream caspase pathway and promote the degradation of the apoptosis-related protein p53 through the ubiquitin-proteasome pathway. Targeting miR-769 with its antagonist to treat cisplatin-resistant GC cells can restore the cisplatin response, confirming that exosomal miR-769-5p can be a key regulator of cisplatin resistance in GC. Therefore, exosomal miR-769-5p derived from drug-resistant cells can be used as a potential therapeutic predictor of anti-tumor chemotherapy to enhance the effect of anti-cancer chemotherapy, which provides a new treatment option for GC.

## Introduction

Gastric cancer (GC) is the leading cause of cancer-related death worldwide [1]. Cisplatin has been widely used for patients with advanced metastatic gastric cancer who are not eligible for surgery [2]. However, not all patients respond to cisplatin, which in turn leads to a poor prognosis [3]. Tumor resistance is a complex dynamic process of mutual influence between individuals and tumors. At the micro-level, it is the result of the mutual adaptation of the tumor microenvironment and tumor cells after chemotherapy [4]. The adaptive changes of tumor cells occur in an orderly manner under the control of intricate signal networks and key molecules, in which the interaction of heredity, epigenetics, and post-translational protein modification has an important role.

MicroRNA (miRNA) is a non-coding RNA with a length of 18-22 nt, which regulates protein expression levels by blocking mRNA translation or inducing mRNA degradation [5]. It can modify the expression of target genes and regulate signal transduction and biological processes [6]. Changes in the expression of certain miRNAs in most tumors have been associated with tumor cell proliferation, angiogenesis, and drug resistance [7, 8].

The apoptotic signaling molecule CASP9 is one of the caspases, a family of proteins that regulates cell death [9, 10]. Anti-apoptosis is an important feature of malignant cells, which has been clearly related to tumor development and cancer resistance to treatment [11]. Targeting anti-apoptosis is considered to be a valuable strategy to improve susceptibility to apoptosis and the response to chemotherapy [12–14].

Another important molecule involved in apoptosis is p53, which can prevent abnormal cell proliferation and canceration and regulation of drug resistance [15, 16]. Evidence shows that up to 80% of cellular proteins are degraded by the ubiquitin-proteasome system (UPS), including p53. UPS is a specialized proteolytic system that controls protein degradation and has an important role in cellular protein homeostasis [17–21].

In this study, we hypothesized that CASP9 and p53 might be a potential target gene of miR-769-5p involved in miR-769-5p’s inhibition of gastric cancer cell apoptosis and might induce cisplatin resistance.

## Results

### miR-769-5p is enriched in BGC823/DDP cell-derived exosomes

To isolate exosomes from BGC823 and BGC823/DDP cells, we purified the conditioned medium by using differential centrifugations. Under the transmission electron microscope, nanovesicles were seen as a round shape with bilayered membranes, and the diameter distribution of these nanovesicles ranged from 40nm to 150 nm for cryopreserved spheres (**Figure 1A**). NanoSight particle tracking analysis (NTA) of the size distributions and a number of exosomes revealed that the size of main vesicles secreted from BGC823 and BGC823/DDP cells was 82 nm and 89 nm, with concentrations of 1.13E+10 particles/ml and 7.29E+9 particles/ml, respectively (**Figure 1B**). By immunoblotting of lysates from purified nanovesicles and flow cytometry (FCM), the known exosomal markers TSG101, CD9, CD81and CD63 were detected (**Figure 1C** and **1D**). These results demonstrated that these nanovesicles isolated from BGC823 and BGC823/DDP express typical characteristics of exosomes.

**Figure. 1.**
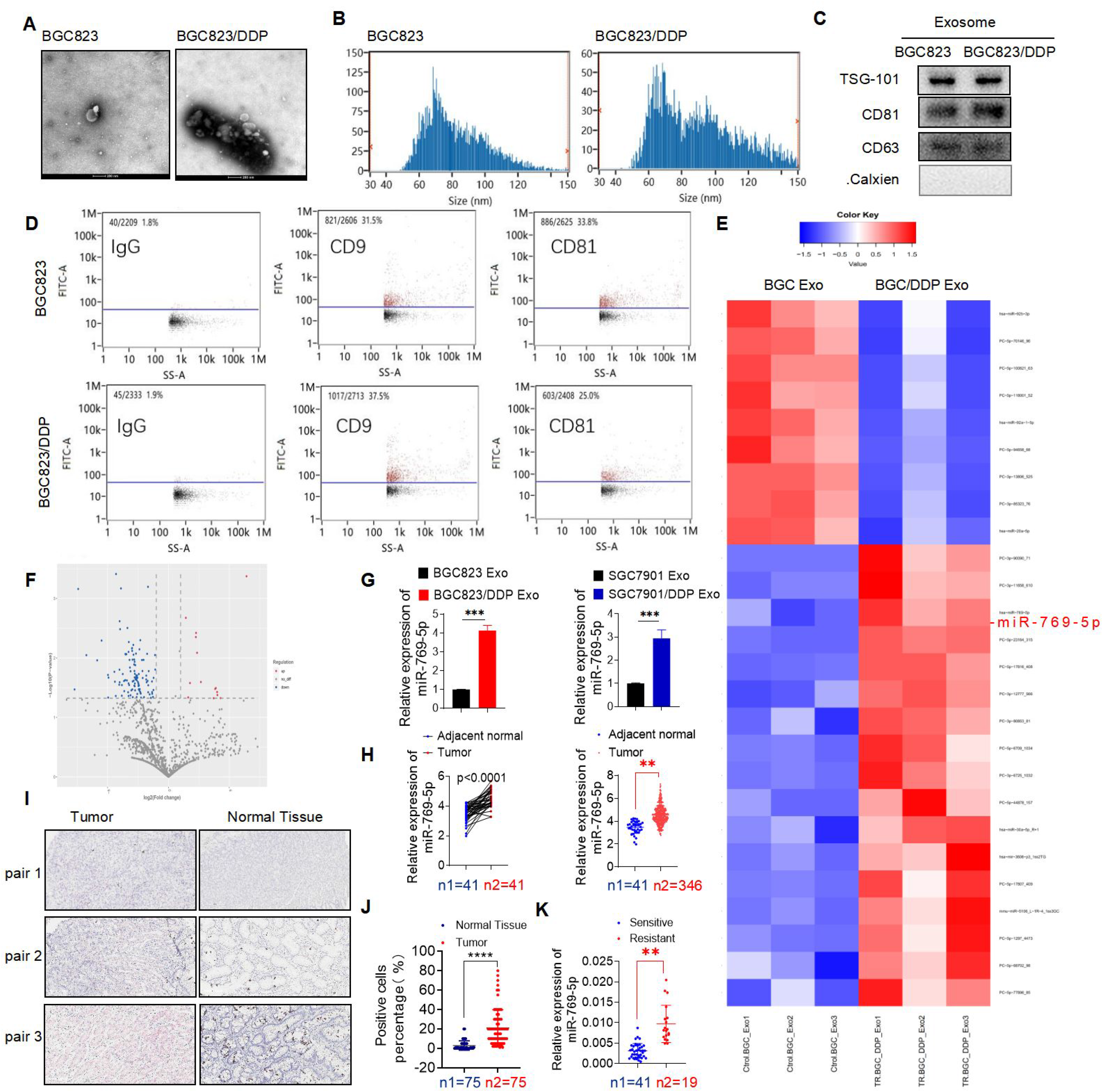
miR-769-5p is enriched in BGC823/DDP cell-derived exosomes. A. Double-membrane exosomes purified from the supernatants of BGC823 and BGC8231/DDP cells were observed by Transmission Electron Microscopy (TEM). B. NanoSight particle tracking analysis (NTA) of the diameter and concentration of vesicles(particles/mL). C, D. Exosomal markers TSG101, CD9, CD81 and CD63 were detected by Western blot and flow cytometry (FCM) to prove that the extract in exosomal protein purified from cell supernatants has the typical characteristics of exosomes. E, F. Cluster heat map and Volcano plot of differential miRNAs in exosomes purified from the supernatants of BGC823 and BGC823/DDP cells. G. qRT-PCR verified the relative expression of miR-769-5p in exosomes purified from the supernatants of BGC823, BGC823/DDP, SGC7901 and SGC7901/DDP cells. H. Different expression of miR-769-5p between 41 pairs of tumor and adjacent tumor, 41 tumors and 346 adjacent tumors according to TCGA database. I, J. The positive rate (referring to the percentage of positive cells) of miR-769-5p in 75 pairs of gastric cancer tissues and adjacent tissues by RNA in situ hybridization (ISH). K. qRT-PCR detected the relative expression of miR-769-5p in serum exosomes of 60 cases (including 41 cisplatin-sensitive cases and 19 cisplatin-resistant cases) of GC patients. After chemotherapy, the level of serum miR-769-5p was significantly increased in non-response patients (n1=19) compared with response patients (n2=41). Quantitative data from three independent experiments are shown as the mean ± SD (error bars). *P < 0.05, **P < 0.01, ***P < 0.001 (Student’s t-test)

Next, we compared the differences in miRNAs expressed in two cell-derived exosome populations by using sequencing analysis (**Figure 1E** and **F**). The level of miR-769-5p expressed in BGC823/DDP secreted exosomes (BD Exo) was 4.77 times that in BGC secreted exosomes (BC Exo). The expression of miR-769-5p in BD Exo was 8.778±0.6923-fold greater than in BC Exo (**Figure 1G**). Moreover, using a TCGA database, we found that miR-769-5p has a promoting role in tumor (**Figure 1H**).

To detect the miR-769-5p expression levels in 79 pairs of clinical samples, we used the technique of RNA in situ hybridization (ISH). Our results revealed that miR-769-5p had markedly higher expression in tumor tissues compared with paracancerous tissues (**Figure 1I** and **1J**). The results indicated that the abundance of miR-769-5p in GC tissues was much higher than that in matched normal tissues, and the expression of miR-769-5p was correlated with advanced TNM stage, vascular invasion and poor prognosis. Additionally, we investigated the expression level of miR-769-5p in human GC serum samples. miR-769-5p expression level was significantly increased in exosomes of DDP-resistant patients’ serum (n=19, as compared to respective parental DDP-sensitive patients’ serum (n=41) (**Figure 1K**). These findings suggested that miR-769-5p may be involved in DDP sensitivity.

### miR-769-5p is is required for GC cisplatin-resistance

Growing evidence indicates that exosomes released by cancer cells are enriched in miRNAs. Exosomal miRNAs can mediate phenotypical changes in the tumor microenvironment (TME) to promote tumor growth and therapy resistance. In this study, we hypothesized that miR-769-5p from BD Exo might participate in this process. We evaluated the effect of DDP on BGC823 cells in the presence of BD Exo and found that BD Exo significantly decreased the sensitivity of BGC823 cells to cisplatin by CCK8 (**Figure 2C**). At a cisplatin concentration of 0.8 ug/ml, the survival of BGC823 cells increased after adding BD Exo compared with control. The half-maximal inhibitory concentration (IC50) of cisplatin was also increased. Additionally, the rates of BGC823 cells’ apoptosis were reduced after being co-cultured with BD Exo for 24h (**Figure 2A** and **2B**). This data suggests that exosomes expression in resistant cells reduces IC50 and increases cell apoptosis of sensitive cells following cisplatin treatment.

**Figure. 2.**
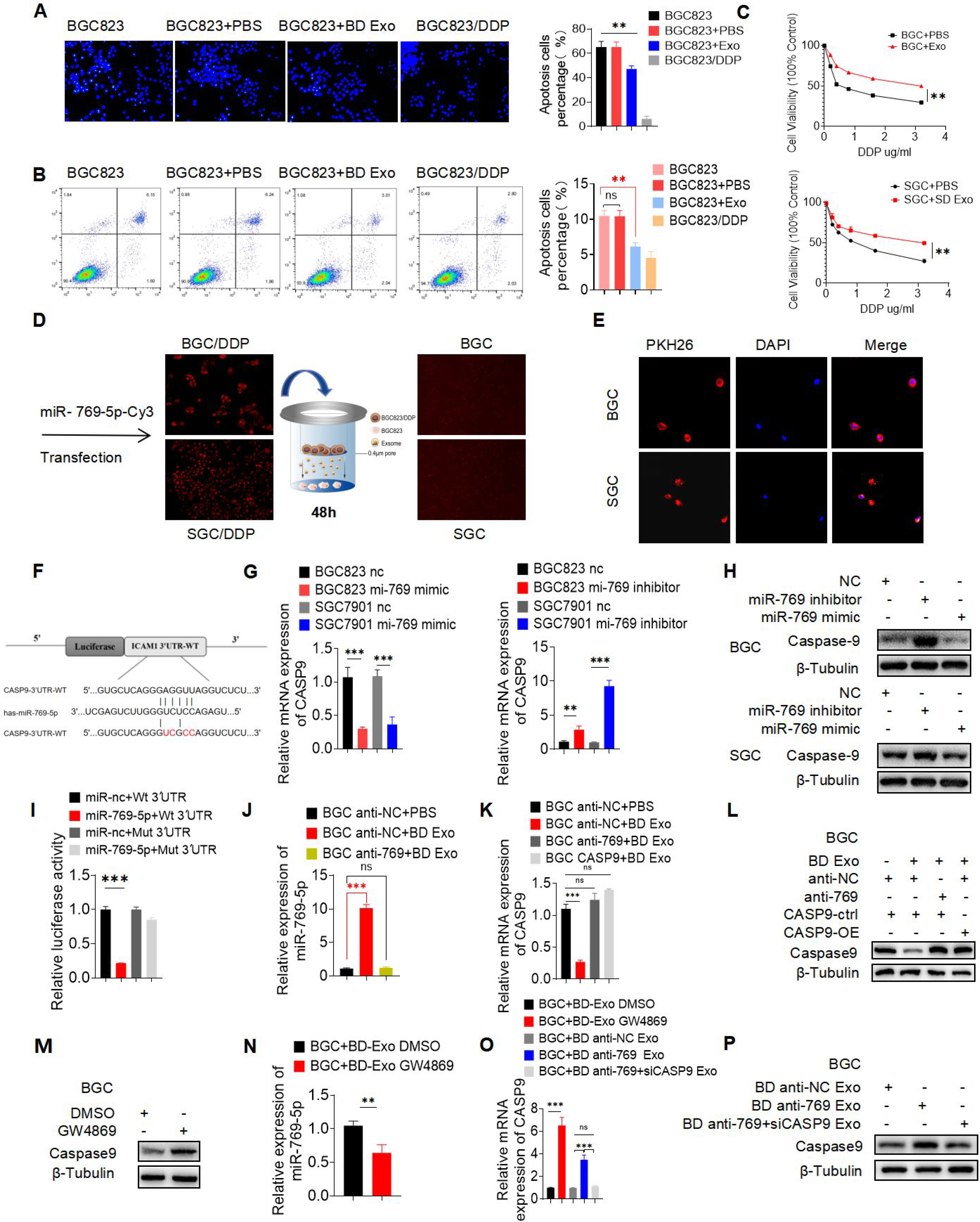
Exosome-mediated transfer of miR-769-5p is required for GC cisplatin-resistance and targets CASP9 directly. A, B. The rates of BGC823 cells’apoptosis were reduced after being co-cultured with BD Exo (200ug/ml) for 24h and treated with cisplatin (0.4 ug/ml) for 24h detected by Hoechst nuclei staining and flow cytometry assay (FCM). C. The survival of BGC823 or SGC7901 cells co-cultured with BD Exo or SD Exo (200ug/ml) for 24h and treated with cisplatin for 24h was detected by CCK-8. D. Red fluorescence was observed in the BGC823 or SGC7901 cells after co-cultured with BGC823/DDP or SGC7901/DDP cells for 24h which were transfected with the Cy3-miR-769-5p mimic (red fluorescence). E. Confocal microscopy showed internalization of exosomes in BGC823 or SGC7901 recipient cells after co-cultured with PKH26-labeled (red fluorescence) BD Exo or SD Exo for 24h. DAPI was used to stain the nuclei of BGC823 or SGC7901 recipient cells with blue fluorescence. F. Predicted binding sites of the CASP9 3′ UTR by miR-769-5p. I. Luciferase reporter was carried out in HEK293T cotransducted with miR-769-5p-mimics or miRNA control with firefly luciferase reporter plasmid containing either wild-type (WT) or mutant (MUT) CASP9 3′ UTR (pGL3-CASP9-WT or pGL3-CASP9-MUT). G, H. PCR and Western blot confirmed that miR-769-5p negatively regulated the expression of CASP9. J. qRT-PCR showed the expression of miR-769-5p in in BGC anti-NC+ PBS, BGC anti-NC + BD Exo and BGC anti-769 + BD Exo. K, L. The mRNA and protein levels of CASP9 in BGC anti-NC+ PBS, BGC anti-NC + BD Exo and BGC anti-769 + BD Exo. M, N. qRT-PCR and Western blot showed the expression of miR-769-5p in BGC+BD Exo DMSO and BGC+BD Exo GW4869. O, P. The upregulation of CASP9 mRNA and protein was detected by qRT-PCR and Western blot in BGC+BD Exo GW4869 and BGC+BD anti-769 Exo. Quantitative data from three independent experiments are shown as the mean ± SD (error bars). *P < 0.05, **P < 0.01, ***P < 0.001 (Student’s t-test)

A Transwell assay was used to examine whether the delivery of miR-769-5p occurs via exosomes. Briefly, we plated BGC823/DDP cells transfected with the Cy3-miR-769-5p mimics in the upper chamber and BGC823 cells in the lower chamber. The co-culture system was separated by 0.4 um pores, just allowing the transmission of micro particles such as exosomes but inhibiting direct contact between cells. After 24h, we found strong red fluorescence in BGC823 cells (**Figure 2D**). This phenomenon proved that miR-769-5p might be directly transferred from donor cells to recipient cells through exosomes. Furthermore, to visualize exosome transfer, we first incubated BGC823 cells and BD Exo in the presence of PKH26-labeled for 24 hours and evaluated the BD Exo uptake levels by measuring the red PKH26 signal in the BGC823 cell line. The confocal immunofluorescence microscopy detected a robust exosome signal in the cytoplasm of BGC823 cells after incubation of labeled BD Exo (**Figure 2E**), thus suggesting that BD Exo was successfully taken up BGC823 cells. **Figure 2J** shows that the co-incubation with BD Exo increased the expression of miR-769-5p. Importantly, intratumor injection of BD Exo promoted the growth and induced the cisplatin resistance of GC cells compared to the same group injected with PBS (**Figure 5D-5G**). Taken together, we have reasons to believe that miR-769-5p might be transferred via exosomes from resistant GC cells to the neighboring sensitive GC cells, thereby spreading cisplatin resistance.

### Exosome-mediated transfer of miR-769-5p targets CASP9 directly

To further explore the mechanism through which BD Exo and miR-769-5p induced cisplatin resistance, we investigated the target gene involved in mediating the effect of miR-769-5p on modulating apoptosis by miRanda, TargetScan, MiRWalk, and miRTarBase. We found that CASP9 was a target of miR769-5p in 3′-UTR area. Luciferase reporter assay further showed a significant reduction in luciferase activity when miR-769-5p was expressed in HEK293T cells as it did not affect the luciferase activity when the binding site was mutated (**Figure 2F** and **2I**). Furthermore, qRT-PCR and Western blotting showed that overexpression of miR-769-5p inhibited the expression of CASP9 in BGC823 cells, whereas inhibition of miR-769-5p reversed this process (**Figure 2G** and **2H**), thus suggesting that miR-769-5p can negatively regulate CASP9 at both the transcript and protein levels.

Next, we infected BGC823 cells with lentiviral vectors to construct cell lines stably expressing miR-769-5p inhibitor (BGC anti-769), negative control miRNA inhibitor (BGC anti-NC), or CASP9 overexpression (BGC CASP9). Then, we cocultured these cells directly with BD Exo (BGC anti-769 + BD Exo, BGC anti-NC + BD Exo and BGC CASP9 + BD Exo), BGC anti-NC incubated with the same amount of PBS (BGC anti-NC+ PBS) were used as a negative control. **Figure 2J** (**SFig 1F**) shows that the co-incubation with BD Exo increased the expression of miR-769-5p in BGC anti-NC but had no effect on the expression of BGC anti-NC in BGC anti-769 cells. Compared with the control group BGC anti-NC+ PBS, the expression of CASP9 in BGC anti-NC + BD Exo was reduced. Nevertheless, when miR-769-5p was inhibited in BGC823, the impact above of reduction in CASP9 induced by BD Exo was offset (**Figure 2K** and **2L**, **SFig 1E** and **1G**). These results suggested that BD Exo can induce the upregulation of miR-769-5p and downregulation of CASP9 in recipient cells.

Transwell assay was used to further explore whether the delivery of miR-769-5p to recipient cells is dependent on exosomes. We plated BGC823/DDP cells with GW4869 in the upper chamber to prevent exocytosis, while BGC823 cells were seeded in the lower chamber. After 24 hours, we collected BGC823 cells and found that the expression of miR-769-5p in the cells (BGC+BD Exo GW4869) was significantly reduced compared with the control group cells treated with DMSO (BGC+BD Exo DMSO)(**Figure 2M**, **SFig 1I**). The CASP9 mRNA and protein expression were significantly increased (**Figure 2N** and **2O**, **SFig 1H** and **1J**). These results indicated that the delivery of miR-769-5p was dependent on exosomes.

In another experiment, we plated BGC823/DDP cells transfected with miR-769-5p inhibitor (BD 769 inhibitor) in the upper chamber and BGC823 cells in the lower one. We found that the co-cultured recipient cells CASP9 mRNA (**Figure 2O**, **SFig 1J**) and protein (**Figure 2P**, **SFig 1K**) levels were higher than the negative control. In addition, when BD cells in the upper chamber were co-transfected with anti-miR-769-5p and CASP9-siRNA (BD anti-769+siCASP9), and exosomes released from BD cells had no statistically significant effect on the mRNA and protein levels of CASP9 in the recipient cells. These results further confirmed that miR-769-5p was present in exosomes and that CASP9 was down-regulated by miR-769-5p.

### Exosome-mediated transfer of miR-769-5p confers cisplatin resistance through downregulating CASP9 and subsequent evasion of apoptosis

Next, we determined that exosomal miR-769-5p confers cisplatin resistance in BGC823 cells by targeting CASP9. As shown in (**Figure 3A, SFig 2A**), BD Exo significantly enhanced the apoptosis of BC anti-NC cells by 4.483 ± 0.3153% induced by cisplatin (0.4 ug/ml, 24h), while no statistically significant difference was observed in BGC823 cells with miR-769-5p knockdown or CASP9 overexpression. Therefore, miR-769-5p knockdown or CASP9 overexpression could reverse the effect of BD Exo on the cisplatin resistance of BGC823 cells. Compared with BGC823/DDP cells treated with DMSO, after co-cultivation with BGC823/DDP cells treated with GW4869 (10 μM), the level of apoptosis of BGC823 cells induced by cisplatin was reduced by 4.470 ± 0.9988% (**Figure 3B**, **SFig 2B**). In addition, when they were co-cultured with miR-769-5p knockdown BGC823/DDP cells, the cisplatin resistance of BGC823 cells was reduced (**Figure 3C**, **SFig 2C**).

**Figure. 3.**
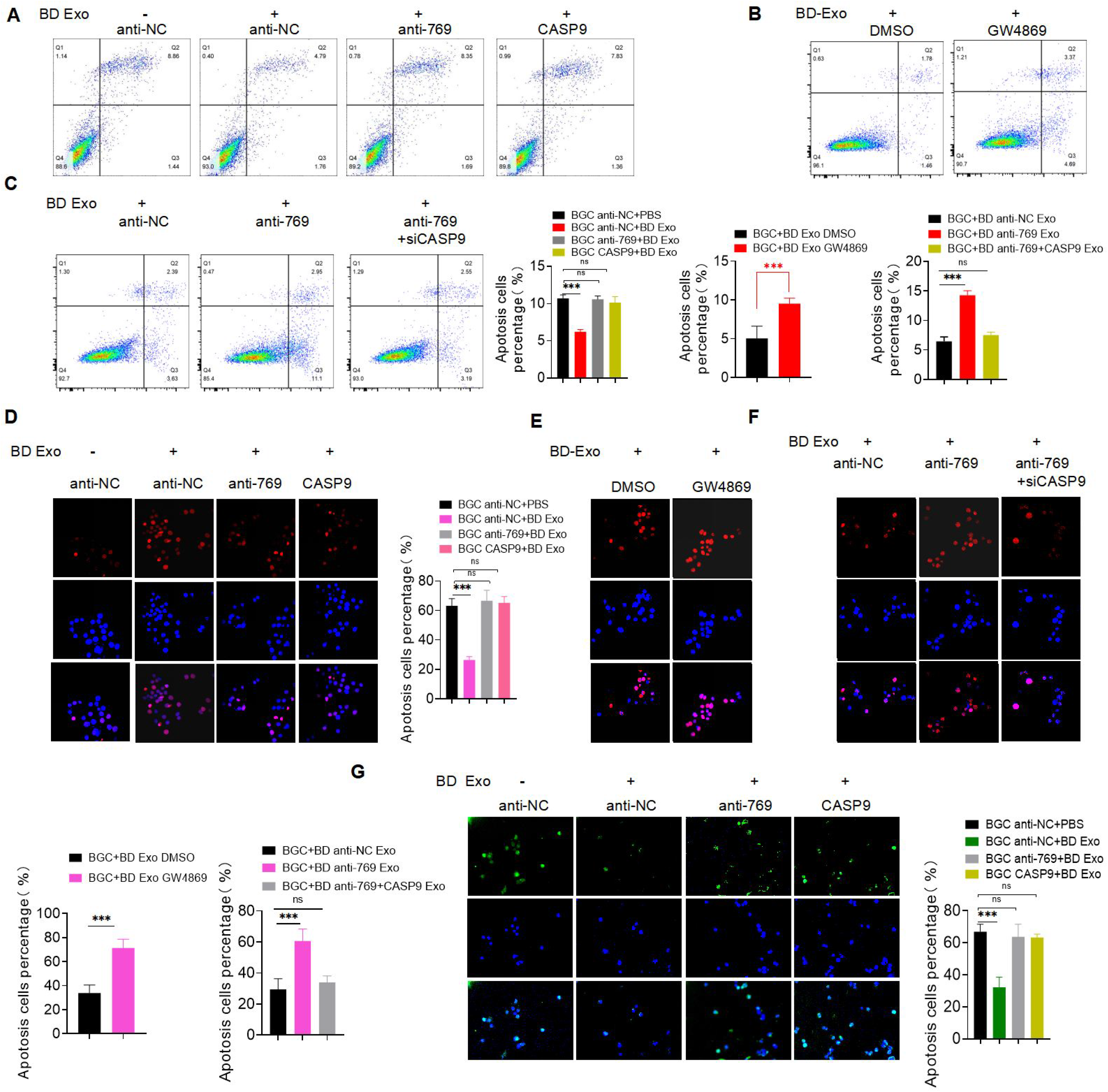
Exosome-mediated transfer of miR-769-5p confers cisplatin resistance through downregulating CASP9. A. Flow cytometry assay detected cell apoptosis rate of BGC anti-NC + PBS, BGC anti-NC + BD Exo, BGC anti-769 + BD Exo and BGC CASP9 + BD Exo. B. Flow cytometry assay detected cell apoptosis rate of BGC+BD Exo DMSO and BGC + BD Exo GW4869. C. Flow cytometry assay detected cell apoptosis rate of BGC + BD anti-NC Exo, BGC + BD anti-769 Exo and BGC + BD anti-769 + siCASP9 Exo. D.The level of γ-H2AX nuclear foci in BGC anti-NC + PBS, BGC anti-NC + BD Exo,BGC anti-769 + BD Exo and BGC CASP9 + BD Exo. E. The level of γ -H2AX nuclear foci in BGC+BD Exo DMSO and BGC+BD Exo GW4869. F. The level of γ-H2AX nuclear foci in BGC+BD anti-NC Exo, BGC + BD anti-769 Exo and BGC+ BD anti-769 + siCASP9 Exo. G. TUNEL analysis detected cell apoptosis rate of BGC anti-NC+PBS, BGC anti-NC + BD Exo, BGC anti-769 + BD Exo and BGC CASP9 + BD Exo. Quantitative data from three independent experiments are shown as the mean ± SD (error bars). *P < 0.05, **P < 0.01, ***P < 0.001 (Student’s t-test)

γ-H2AX is a sign of DNA double-strand breaks. After 24 h cisplatin treatment (0.8 ug/ml, 24h), the level of γ-H2AX nuclear foci in the control group remained high, but the nuclear foci in the BD Exo co-culture group significantly decreased by 36.77 ± 3.079% (**Figure 3D**, **SFig 3A**). However, there was no statistically significant difference observed in BGC823 cells with miR-769-5p knockdown or CASP9 overexpression. γ-H2AX expression levels in nuclear foci indicated that cisplatin induces more resistant cell lines after co-culturing with BD Exo. Similarly, after co-culturing with BGC823/DDP cells treated with GW4869 (10 μM), the level of γ-H2AX expression in nuclear foci of BGC823 cells induced by cisplatin was reduced by 37.47 ± 5.590% compared with BGC823/DDP cells treated with DMSO (**Figure 3E**, **SFig 3B**).

We then used a Transwell assay and co-cultured BGC823/DDP cells transfected with miR-769-5p inhibitor (BD anti-769) with BGC823 cells seeded in the lower chamber and found that γ-H2AX expression levels in nuclear foci of co-cultured recipient cells were higher than the negative control (**Figure 3F**, **SFig 3C**). Co-incubation of BGC823/DDP cells co-transfected with miR-769-5p inhibitor and CASP9-siRNA had no profound synergistic effect on γ-H2AX expression in BGC823 cells.

To further investigate the role of exosomal miR-769-5p cisplatin-induced apoptosis, we performed TUNEL analysis and found that it was consistent with the verification of flow cytometry assays (**Figure 3G**, **Figure 4A** and **B**, **SFig 3D-3F**).

**Figure. 4.**
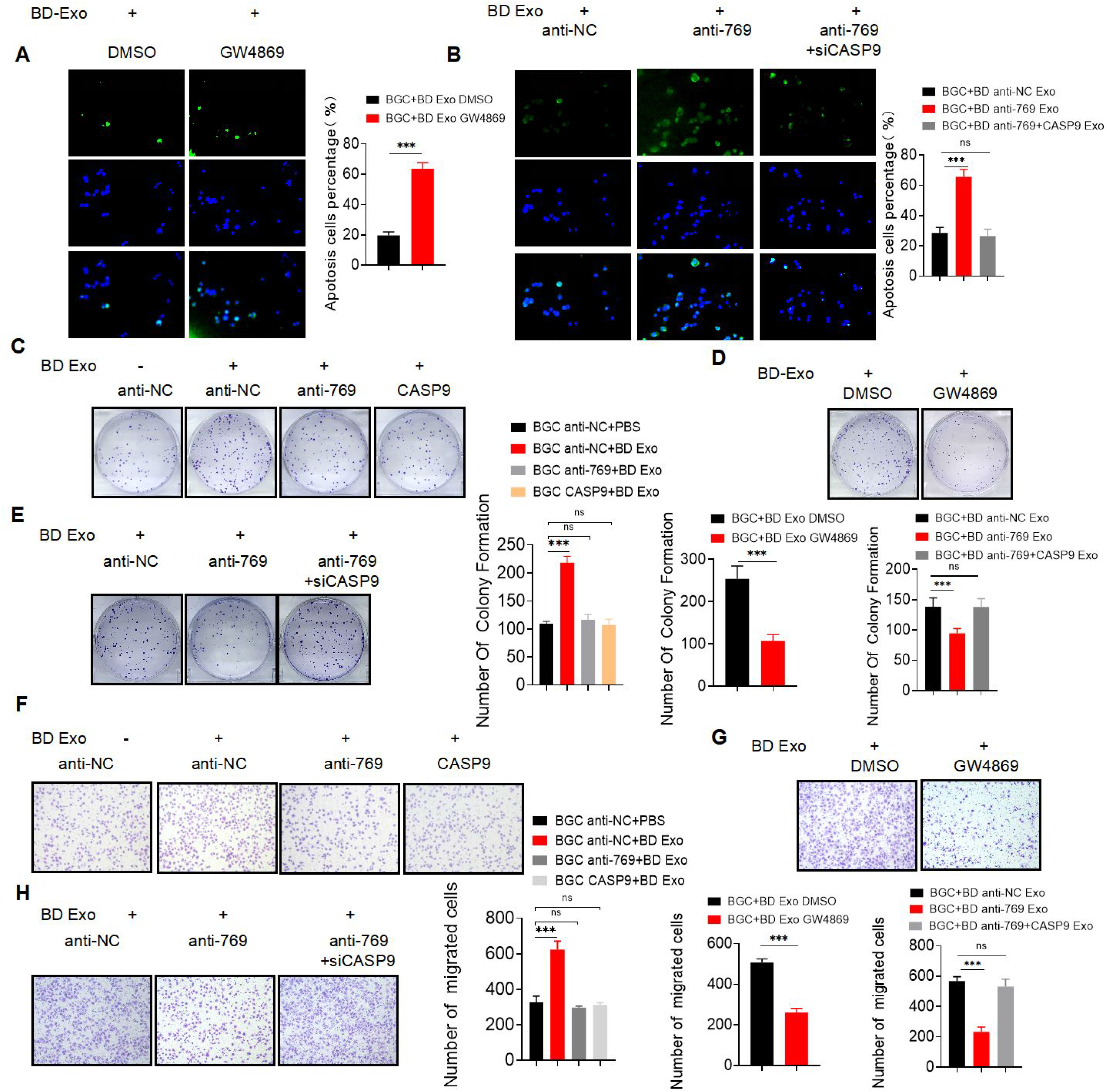
Exosomal miR-769-5p promotes recipient cells proliferation and migration by downregulating CASP9. A. TUNEL analysis detected cell apoptosis rate of BGC + BD Exo DMSO and BGC + BD Exo GW4869. B. TUNEL analysis detected cell apoptosis rate of BGC + BD anti-NC Exo, BGC + BD anti-769 Exo and BGC + BD anti-769 + siCASP9 Exo. C. The average colony numbers of three independent experiments were calculated in BGC anti-NC + PBS, BGC anti-NC + BD Exo, BGC anti-769 + BD Exo and BGC CASP9 + BD Exo. D. The average colony numbers of three independent experiments were calculated in BGC + BD Exo DMSO and BGC + BD Exo GW4869. E. The average colony numbers of three independent experiments were calculated in BGC + BD anti-NC Exo, BGC + BD anti-769 Exo and BGC + BD anti-769 + siCASP9 Exo. F. Migration ability of BGC anti-NC + PBS, BGC anti-NC + BD Exo, BGC anti-769 + BD Exo and BGC CASP9 + BD Exo were assessed by Transwell assay. G. Migration ability of BGC + BD Exo DMSO and BGC+BD Exo GW4869 were assessed by Transwell assay. H. Migration ability of BGC+BD anti-NC Exo, BGC + BD anti-769 Exo and BGC + BD anti-769 + siCASP9 Exo were assessed by Transwell assay. Quantitative data from three independent experiments are shown as the mean ± SD (error bars). *P < 0.05, **P < 0.01, ***P < 0.001 (Student’s t-test)

The results showed that the exosomal miR-769-5p from cisplatin-resistant cells could accelerate cell apoptosis of cisplatin-sensitive cells. Western blots assay demonstrated that the protein levels of caspase-9 and cleaved caspase-3 in BGC anti-NC + BD Exo cells were reduced, yet there were no obvious differences in the BGC anti-769 + BD Exo and BGC CASP9 + BD Exo cells (**Figure 5A**, **SFig 4A**). Compared with BGC+BD Exo DMSO or BGC+BD anti-NC Exo cells, the caspase-9 and cleaved caspase-3 protein levels were increased in BGC823 cells co-cultured with BGC823/DDP cells treated with GW4869 or transfected with miR-769-5p inhibitor (**Figure 5B** and **5C**, **SFig 4B** and **4C**). Thus, these data suggested that the knockdown miR-769-5p could reverse the chemoresistance of gastric cancer cells to cisplatin.

**Figure. 5.**
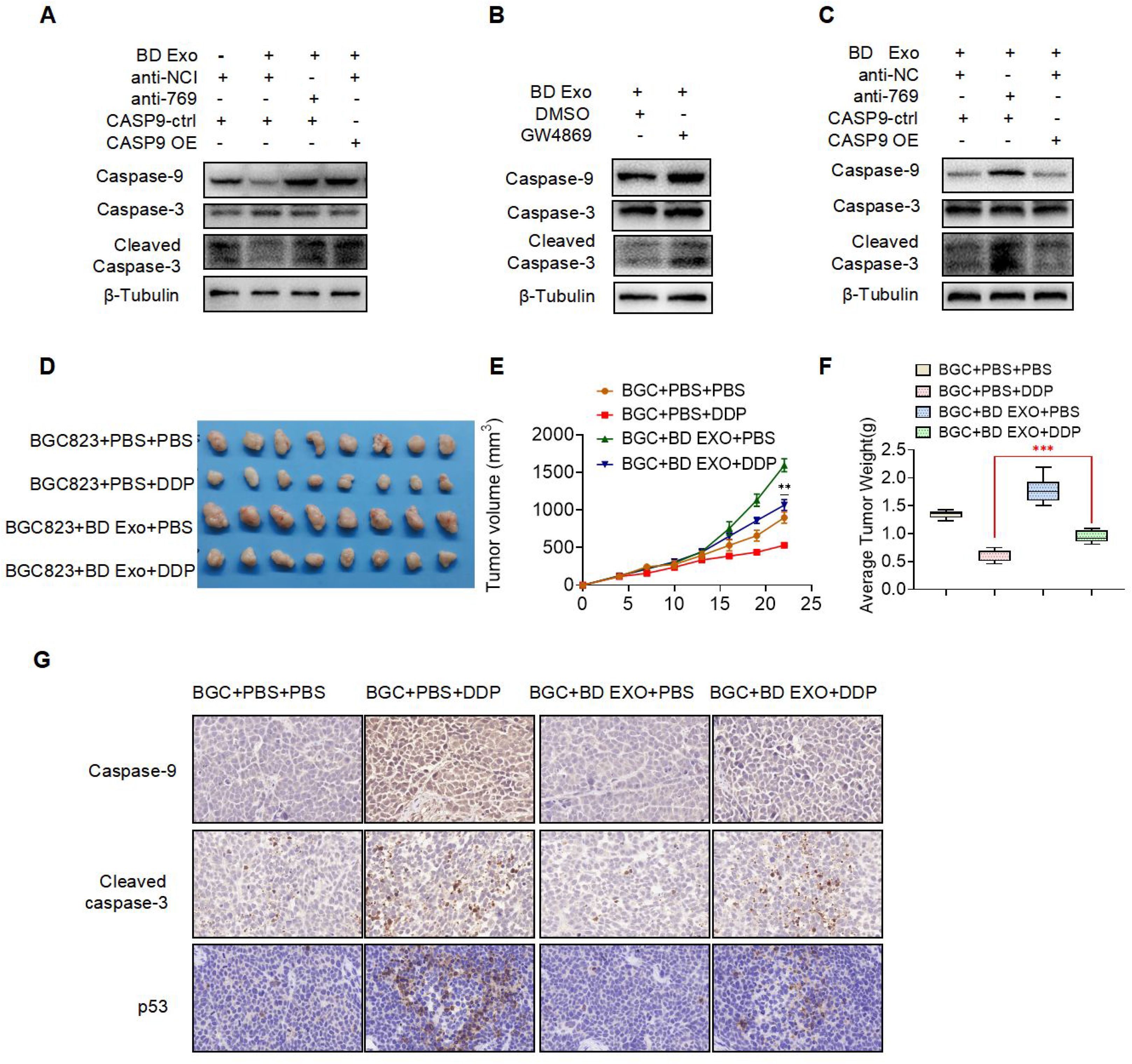
Exosomal miR-769-5p confers cisplatin resistance through downregulating CASP9 along with subsequent evasion of apoptosis and confirmed in vivo. Western blot anaysis of caspase9, caspase3 and cleaved caspase3 in BGC + BD anti-NC Exo, BGC + BD anti-769 Exo and BGC + BD anti-769 + siCASP9 Exo. B. Western blot anaysis of caspase9, caspase3 and cleaved caspase3 in BGC + BD Exo DMSO and BGC + BD Exo GW4869. C. Western blot anaysis of caspase9, caspase3 and cleaved caspase3 in BGC + BD anti-NC Exo, BGC + BD anti-769 Exo and BGC + BD anti-769 + siCASP9 Exo. D. Subcutaneous xenograft assay of BGC823 cells with or without BD Exo (200ug/100μL cells per mouse) once every two days in nude mice with PBS or cisplatin (DDP, 4mg/kg) treatment. E. Tumor volume of xenograft models were measured every two days and shown. *Tumor volume (mm^3^) = 0.5 ×width^2^ × length.* F. Tumor weight of xenograft models were measured every two days and shown. G. CASP9, cleaved caspase3 and p53 expression levels were shown in representative xenograft tumors by Immunohistochemistry (IHC) (400x magnification, scale bars = 50 μm). Results are presented as mean SD. *P < 0.05, **P < 0.01, ***P < 0.001

### Exosomal miR-769-5p promotes recipient cells proliferation and migration by downregulating CASP9

Next, we investigated whether exosomal miR-769-5p affects the biological processes of GC cells. BGC anti-NC cells treated with BD EXO showed increased colony formation, migration capacity compared to BGC anti-NC cells treated with PBS (**Figure 4C** and **4F**, **SFig 2D** and **2G**). Nevertheless, this alteration was reversed when BGC anti-769 or BGC CASP9 cells were co-cultured with BD Exo. In contrast, when BGC823 cells were co-cultured with BGC823/DDP treated with GW4869 or miR-769-5p knockdown, the colony formation, migration capacity of BGC823 cells decreased compared to those of the corresponding negative controls (Figure 4D and 4E, 4G and 4H, **SFig 2E** and **2F**, **SFig 2H** and **2I**). Our findings suggested that exosomal miR-769-5p enhanced GC cell proliferation and migration by downregulating CASP9.

To sum up, the miR-769-5p was markedly upregulated in BGC823 cells treated with BD Exo, which suggested its potential role in cisplatin resistance and indicated the possibility of achieving the cisplatin resistance through the exosomal transfer of miR-769-5p by inhibiting CASP9 in GC cells.

### miR-769-5p promotes ubiquitin-mediated p53 protein degradation in GC cells

It has been reported that the transcription factor P53 is essential in the complex molecular network regulating apoptosis, and the activation of tumor suppressor P53 is crucial for preventing abnormal cell proliferation and carcinogenesis. Many studies have shown that P53 is involved in regulating the generation of drug resistance. The main targets of P53 include P21, PUMA, BAX, and BID [22–24]. To further determine whether miR-769-5p is involved in GC cisplatin resistance and its molecular mechanism, we found that the targets of differentially expressed miRNAs were enriched in the p53 pathway based on the KEGG enrichment analysis of differently expressed miRNAs in exosomes (**Figure 6A**). Therefore, we hypothesized that miR-769-5p might affect the p53 pathway. To evaluate whether miR-769-5p is involved in p53-mediated apoptosis of gastric cancer cells, miR-769-5p expression in BGC823 and SGC7901 cells was overexpressed and knocked down using miR-769-5p mimics and inhibitors, respectively, after which the expression of p53 mRNA and protein were analyzed (**Figure 6B** and **6C**). Western blotting showed that miR-769-5p silencing significantly enhanced the expression of p53 in GC cells, while overexpression of miR-769-5p had the opposite effects (**Figure 6C**). It indicated that miR-769-5p negatively regulates p53 protein expression and p53-mediated apoptosis in gastric cancer cells. However, qRT-PCR showed that the transcription level of p53 was not affected by miR-769-5p in gastric cancer cells, indicating that the p53 protein in gastric cancer cells may be degraded by ubiquitination (**Figure 6B**). As a result, we transfected miR-769-5p inhibitors into GC cells, and twenty-four hours later, the cells were treated with 20 μg/ml cycloheximide (CHX) changes with treatment time (0h, 1h, 4h). The cell lysates were then collected within a specified time period and analyzed by Western blot. Higher expression of p53 protein was detected in the cells treated with CHX compared with negative controls (**Figure 6E**, **SFig 4D**). We also treated the cells with MG-132, a specific inhibitor of a ubiquitin-binding protein, and found that higher expression of p53 protein was detected in the cells treated with MG-132 (10um) for 6h (**Figure 6F**, **SFig 4E**), indicating that p53 protein degradation depends on the ubiquitination.

**Figure. 6.**
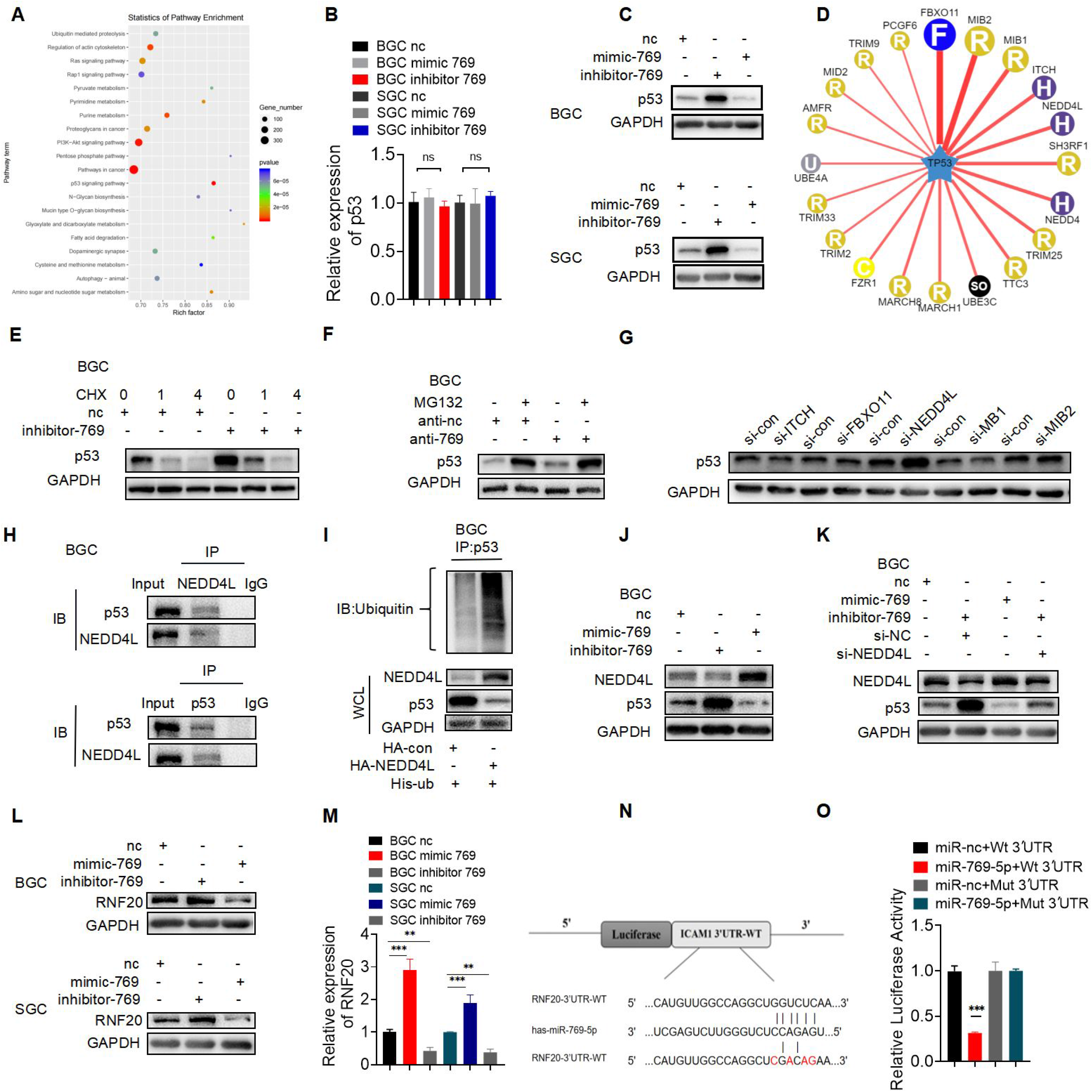
miR-769-5p promotes ubiquitin-mediated p53 protein degradation in GC cells. A. KEGG enrichment analysis showed that the target genes of differentially expressed miRNAs are enriched in the p53 pathway. B. qPCR detected the expression level of p53 mRNA in BGC NC, BGC mimic-769 and BGC inhibitor-769. C. Western blot analysis of expression level of p53 protein in in BGC NC, BGC mimic-769 and BGC inhibitor-769. D. UbiBrowser website predicted E3 ubiquitination ligase with p53 as a substrate. E. Western blot analysis of p53 protein level of 100ug/ml treated with cycloheximide (CHX) changes with treatment time (0h, 1h, 4h). F. Analysis of p53 protein level by Western blot in BGC nc and BGC inhibitor-769 after treatment of MG-132 (10um) for 6h. G. Western blot analysis of p53 protein expression level after transfection of E3 ubiquitinated ligase specific small interfering RNA (siRNA): siFBXO11, siMIB2, siMIB1, siITCH and siNEDD4L. H. Co-IP detected the interaction between NEDD4L and p53 in gastric cancer cells. I. Co-IP and western blot detected p53 ubiquitination modification mediated by NEDD4L. J, K. The expression of NEDD4L and p53 protein levels when miR-769-5p is knocked down or overexpressed. L, M. PCR and Western blot verified the negatively regulatory effects of miR-769-5p on RNF20. N. Predicted binding sites of the RNF20 3′ UTR by miR-769-5p. O. Luciferase reporter was carried out in HEK293T cotransducted with miR-769-5p-mimics or miRNA control with pGL3-RNF20-WT or pGL3-RNF20-MUT. Quantitative data from three independent experiments are shown as the mean ± SD (error bars). *P < 0.05, **P < 0.01, ***P < 0.001 (Student’s t-test)

According to ubibrowser, we characterized the p53-specific E3 ubiquitin ligases to determine the mechanism of miR-769-5p mediated p53 ubiquitination in GC cells. We selected the top five p53 E3 ubiquitin ligases to be silenced by sequence-specific small interfering RNA (siRNA) in HEK-293T. Detection of p53 protein showed that when NEDD4L expression is knocked down by sequence-specific siRNA, p53 levels increase (**Figure 6G**). NEDD4L is the key E3 ubiquitin ligase for p53 ubiquitination in GC cells [25–27]. However, the negative control of NEDD4L-siRNA did not affect p53 expression.

Co-immunoprecipitation (Co-IP) and Western blotting were used to detect the interaction between NEDD4L and p53 in gastric cancer cells (**Figure 6H** and **6J, SFig 4F** and **SFig 4H**). The NEDD4L overexpression plasmid and His-Ub plasmid were co-transfected in BGC, and the ubiquitination level of p53 was detected by immunoprecipitation and Western blotting. NEDD4L overexpression promoted the ubiquitination of p53 (**Figure 6I**, **SFig 4G**), indicating that NEDD4L mediates the ubiquitination modification. In order to further evaluate the effect of miR-769-5p on the expression of NEDD4L, we inhibited and overexpressed miR-769-5p in gastric cancer cell lines to detect the expression of NEDD4L and p53 protein levels (**Figure 6K**, **SFig 4I**). Compared with the negative control group, knockdown of miR-769-5p significantly reduced the expression of NEDD4L and increased the expression level of p53, while overexpression of miR-769-5p showed the opposite result. Western blot also demonstrated that NEDD4L silencing caused p53 protein accumulation in miR-769-5p-silenced cancerous cells. This indicated that the inhibition of miR-769-5p could inhibit the expression of E3 ubiquitinated ligase NEDD4L, increasing the level of substrate p53. These data suggested that miR-769-5p could promote NEDD4L’s expression, leading to its participation in the p53 ubiquitination degradation process.

### E3 ubiquitination ligase RNF20 participates in miR-769-5p mediated p53 protein ubiquitination in GC cells

According to the miRNAs target gene prediction, we found that NEDD4L was not the target gene of miR-769-5p. So, it was unclear how miR-769-5p regulates and inhibits the expression of NEDD4L. Based on the miRNA target gene prediction website and UbiBrowser website, we found that E3 ubiquitin ligase RNF20 might be the target gene of miR-769-5p (**Figure 6L**, **6M** and **6N**). To characterize the interaction between miR-769-5p and RNF20, a dual-luciferase reporter assay was conducted in HEK293T cells. The results revealed that miR-769-5p significantly decreased the activity of the reporter luciferase that was fused with the wild-type RNF20 3-untranslated region (UTR) compared with the controls (**Figure 6O**). This observation suggested a direct interaction between miR-769-5p and RNF20 mRNA. Reports showed that a low RNF20 level was correlated with shortened overall survival and disease-free survival, indicating poor prognosis in cancers [28, 29].

Additionally, we discovered that RNF20 and NEDD4L interacted in GC cells. We transfected silenced and overexpressed RNF144B and negative control plasmids in BGC823 and tested the effect of RNF20 on apoptosis by TUNEL experiment (**Figure 7A** and **7D**, **SFig 5A** and **5C**) and immunofluorescence detection of γ-H2AX expression level (**Figure 7B** and **7E**, **SFig 5B** and **5D**). TUNEL results showed that compared with the negative control group, overexpression of RNF20 significantly promoted the apoptosis of gastric cancer cells while inhibition of RNF20 inhibited cell apoptosis. The results of immunofluorescence detection of γ-H2AX expression level were consistent with the results of the TUNEL experiment. Moreover, Western blot showed that the overexpression of RNF144B resulted in increased cleaved caspase 3 and related to activated apoptosis (**Figure 7C** and **7F**, **SFig 6A** and **6B**). The activation of apoptosis by RNF20 overexpression was further confirmed by flow cytometry assay (**Figure 7H**). The above results indicate that RNF20, as a target gene of miR-769-5p, can participate in cell apoptosis.

**Figure. 7.**
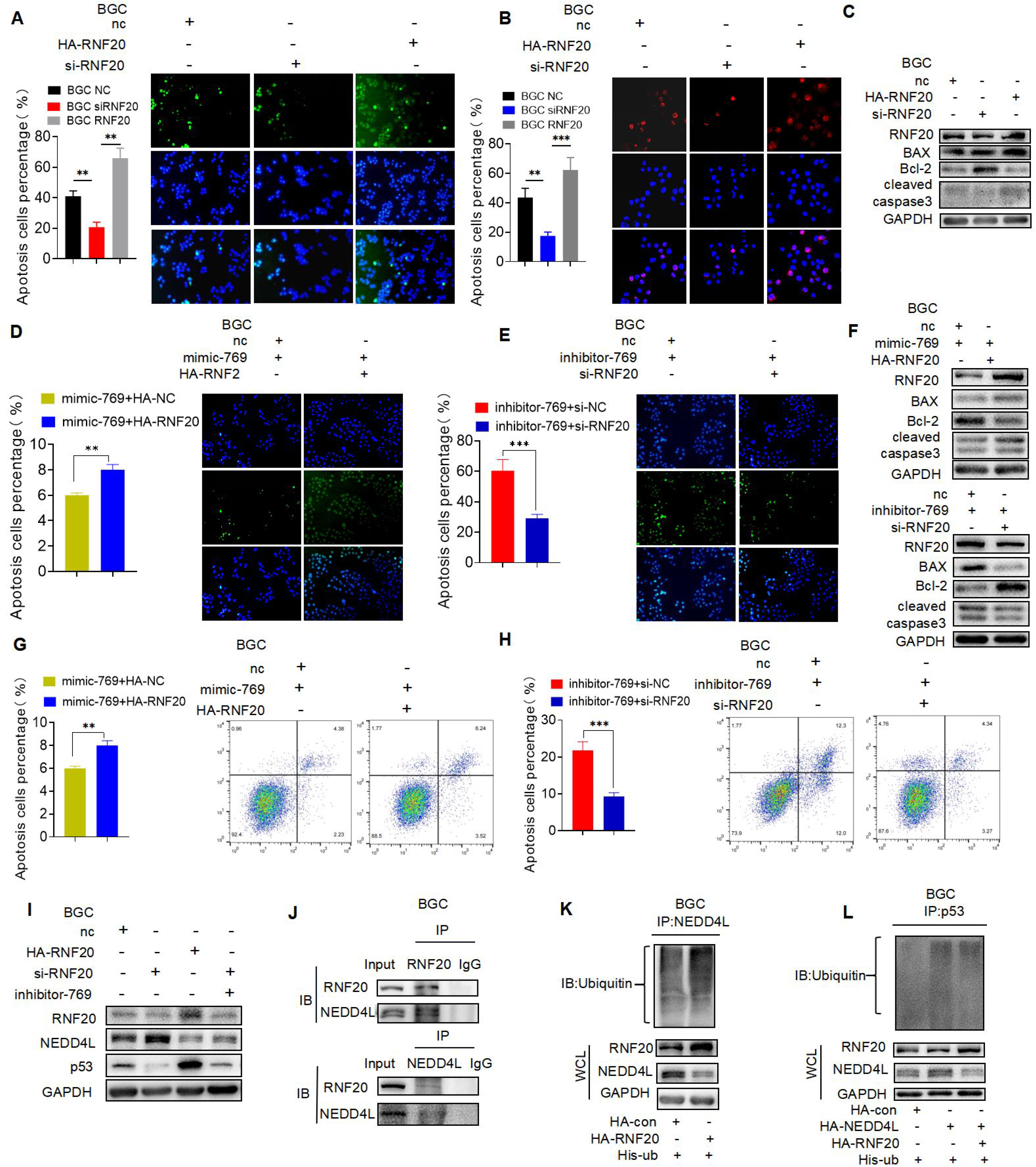
E3 ubiquitination ligase RNF20 participates in miR-769-5p mediated p53 protein ubiquitination in GC cells. A. TUNEL analysis detected cell apoptosis rate of BGC NC, BGC HA-RNF20 and BGC si-RNF20. B. The level of γ -H2AX nuclear foci in BGC NC, BGC HA-RNF20 and BGC si-RNF20. C. The western blot analysis of Bax, Bcl-2 and cleaved caspase 3 proved the positive mediation of RNF20 on apoptosis. D, E, F. The recovery proved that miR-769-5p inhibits the process of apoptosis by down-regulating RNF20 by analysis of TUNEL and western blot. G, H. Flow cytometry assay proved that miR-769-5p inhibits the process of apoptosis by down-regulating RNF20. I. The protein levels of NEDD4L and p53 when RNF20 overexpression and knockdown. J. Co-immunoprecipitation proves that NEDD4L interacts with RNF20. K, L. Co-immunoprecipitation proves that the ubiquitination modification of NEDD4L is mediated by RNF20 Quantitative data from three independent experiments are shown as the mean ± SD (error bars). *P < 0.05, **P < 0.01, ***P < 0.001 (Student’s t-test)

RNF20 can be used as a target gene of miR-769-5p to participate in cell apoptosis. Thus, we further determined how RNF20 conveys apoptotic signals in p53-mediated cell apoptosis. The gene expression of RNF20 was silenced or overexpressed in GC cells, followed by RNF20 and p53 protein detection. RNF20 overexpression markedly suppressed NEDD4L expression and simultaneously induced p53 expression in gastric cancer cells (**Figure 7G**, **SFig 6C**), while silencing the RNF20 gene had the opposite effect on NEDD4L and p53 expression in gastric cancer cells. Furthermore, we overexpressed the RNF20 plasmid in GC cells and performed Co-IP with anti-RNF20 to identify proteins that interacted with RNF20. Our results indicated that RNF20 was bound to NEDD4L (**Figure 7I**, **SFig 6D**), thus suggesting that RNF20 participates in p53-mediated gastric cancer cell apoptosis by regulating NEDD4L expression.

To clarify whether NEDD4L could be ubiquitinated by RNF20 (**Figure 7Q** and **7R**, **SFig 6E** and **6F**), the His-Ub and RNF20 were co-expressed in GC cells, and anti-NEDD4L were used to pull down modified proteins. The presence of polyubiquitinated NEDD4L was observed as a smeared band because of the heterogeneous modification of this protein. At the same time, we stained the polyubiquitinated NEDD4L in the flag-Ub immunoprecipitants to confirm that the ubiquitination modification of NEDD4L was mediated by RNF20 and found that RNF20 overexpression further enhanced the polyubiquitinated NEDD4L compared with the control. These findings revealed that RNF20 was an E3 ligase for NEDD4L and that RNF20 polyubiquitinated NEDD4L for degradation.

### Exosomal miR-769-5p induces cisplatin resistance and promotes the tumorigenesis of GC *in vivo*

Given the observed effects of exosomal miR-769-5p on GC cells *in vitro*, we subsequently confirmed the aforementioned results *in vivo*. To determine whether miR-769-5p sensitizes GC cells to chemotherapeutic agents *in vivo*, anti-miR-769-5p transfected BGC823/DDP cells were subcutaneously implanted into nude mice and then treated with cisplatin (DDP, 4mg/kg). Our data indicated that miR-769-5p knockdown significantly decreased cisplatin resistance in BGC823/DDP xenografts (**Figure 8A-8E**). Levels of exosomal miR-769-5p were approximate two folds lower in the serum than that of the negative control group, and the expression levels of CASP9, p53, and cleaved caspase3 were decreased when the level of miR-769-5p increased in the subcutaneous tumor tissues of mice (**Figure 8N**). These data support our findings that knockdown miR-769-5p ameliorates cisplatin-resistant GC *in vitro* and *in vivo*.

**Figure. 8.**
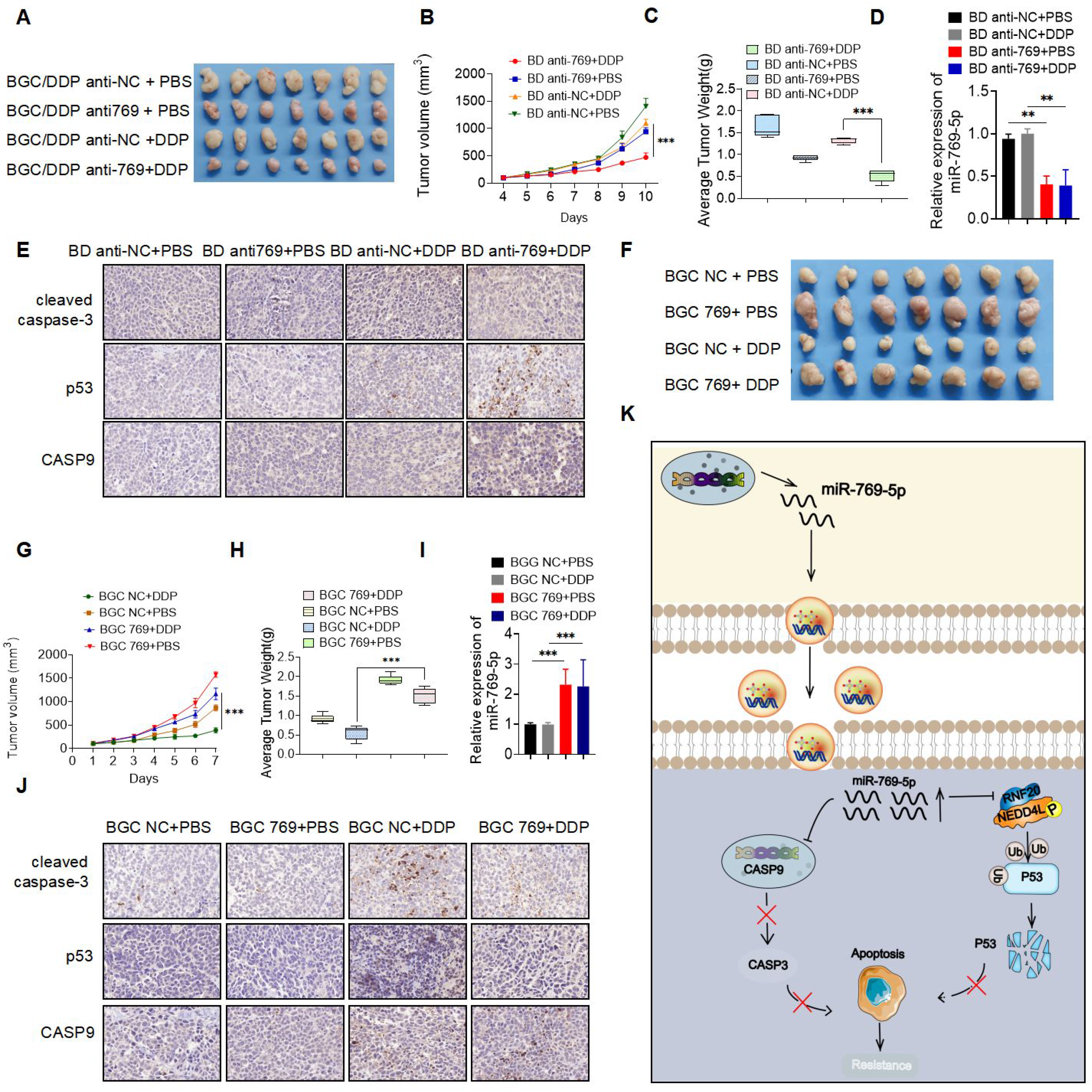
Exosomal miR-769-5p induces cisplatin resistance and promotes the tumorigenesis of GC in vivo. A. Subcutaneous xenograft assay of BGC823/DDP cells (5 × 10^6^ cells/100μL) with or without miR-769-5p knockdwon in nude mice with PBS or cisplatin (DDP, 4mg/kg) treatment. B. Tumor volume of xenograft models were measured every two days and shown. C. Tumor weight of xenograft models were measured every two days and shown. D. Levels of exosomal miR-769-5p in the serum were detected by qPCR. E. CASP9, cleaved caspase3 and p53 expression levels were shown in representative xenograft tumors by Immunohistochemistry (IHC) (400x magnification, scale bars = 50 μ m). F. Subcutaneous xenograft assay of BGC823 cells (5 × 10^6^ cells/100μL) with or without miR-769-5p overexpressed in nude mice with PBS or cisplatin (DDP, 4mg/kg) treatment. G. Tumor volume of xenograft models were measured every two days and shown. H. Tumor weight of xenograft models were measured every two days and shown. I. Levels of exosomal miR-769-5p in the serum were detected by qPCR. J. CASP9, cleaved caspase3 and p53 expression levels were shown in representative xenograft tumors by Immunohistochemistry (IHC) (400x magnification, scale bars = 50 μ m). K. Summary of the mechanism by which exosomal miR-769-5p induces cisplatin resistance. Results are presented as mean SD. *P < 0.05, **P < 0.01, ***P < 0.001

In addition, we subcutaneously injected the stably transfected BGC 823 NC and BGC823 769 cells into nude mice and found that the tumors of BGC823 769 grew faster than those of BGC823 NC. After cisplatin treatment, the tumor volume of the BGC823 769 group was significantly higher compared to BGC 823 NC group (**Figure 8F-8J**). These results indicated that miR-769-5p could promote growth and induce the cisplatin resistance of BGC823 cells *in vivo*. Collectively, miR-769-5p expression was indispensable for cisplatin resistance in GC cells.

## Discussion

Chemotherapy is the most important treatment for patients who cannot undergo surgery or those with advanced metastatic gastric cancer [30]. Yet, multidrug resistance, which has been associated with a poor prognosis, remains a big challenge when treating cancer patients [31]. For example, cisplatin resistance presents a big obstacle in treating patients with advanced gastric cancer.

miRNAs can be encapsulated in exosomes to avoid degradation. Exosomal miRNA can be transported to recipient cells and change their phenotype through changes in gene expression [32–34]. For example, drug-resistant cancer cells may release exosomal miRNAs into the microenvironment, causing the recipient cells to develop drug resistance [35–37]. This ability of exosomes shed from tumor-resistant cells to transfer drug-resistant phenotypes to drug-sensitive cells is considered an important mechanism of drug resistance that is mainly spread mainly by drug efflux pump and miRNAs’ transfer. Numerous studies have reported that exosomes have an important role in invasive tumor progress and chemotherapy resistance.

Our results showed that miR-769-5p in exosomes derived from cisplatin-resistant cells could confer drug-resistant phenotypes on recipient cells and alter their gene expression and apoptosis. This is because when BGC823 cells treated with exosomes respond to cisplatin, the survival time increases. We also found that BD Exo inhibits the effect of cisplatin in BGC823 cells by transferring miR-769-5p. However, transfection of anti-miR-769 into BD cells partially blocked the effect of BD exo on cisplatin. **Figure 8k** summarizes the mechanism through which drug-resistant cells transfer mir-769-5p-loaded exosomes to sensitive cells and modulated cisplatin resistance. Mechanistically, exosomal miR-769-5p inhibits cell apoptosis by targeting the downstream caspase pathway of CASP9 inactivation and enhancing the drug resistance of recipient cells to cisplatin **(Fig. 8K)**.

The activation of the tumor suppressor p53 is essential to prevent abnormal cell proliferation and canceration. Many studies have shown that p53 is involved in the regulation of drug resistance. For example, phosphorylation of p53 serine 15 (Ser15) and serine 20 (Ser20) has been identified as essential in cisplatin resistance [38, 39]. As a key cellular protein regulator, ubiquitination can cause protein degradation. In the process of protein ubiquitination, E3 ubiquitin ligase determines substrate specificity and substrate selection. In addition, the mechanism of ubiquitin-mediated p53 protein degradation has been extensively studied [40, 41]. For example, mdm2-dependent p53 polyubiquitination and degradation can regulate cell proliferation, DNA damage-induced apoptosis, and tumorigenesis by inhibiting p53 [42, 43]. However, the role of miRNA in the regulation of p53 protein ubiquitination remains unclear.

Looking for the target genes of miR-769-5p, we found that miR-769-5p promotes the degradation of p53 and inhibits apoptosis through the ubiquitin-proteasome pathway, thus promoting the resistance of gastric cancer cells to cisplatin. Our study revealed a new mechanism of p53 protein ubiquitination mediated by miR-769-5p in cisplatin resistance. As an important apoptosis-related protein, miR-769-5p participates in the apoptosis of gastric cancer cells through the RNF20-NEDD4L-p53 pathway in the process of induced apoptosis, and miR-769-5p can directly inhibit the expression of RNF20. Previous studies have shown that HBRE1 /RNF20 is the E3 ubiquitin ligase of hiprotein H2B, and the deletion of RNF20 as a tumor suppressor can lead to the overall decrease of H2Bub level[44, 45]. Our results showed that RNF20 had a critical role in p53 protein ubiquitination in gastric cancer cells, mediating the direct degradation of p53 protein by E3 ubiquitin ligase NEDD4L, thus revealing a novel miRNA-mediated p53 protein ubiquitination pathway (**Figure 8K**).

Cancer is a complex genetic disease. Chemotherapy and radiation therapy have always been the core treatment options for cancer. However, these treatments have adverse side effects. Due to malignant tumors being highly heterogeneous in their occurrence and development, this study proved that miR-769-5p, which is highly expressed in drug-resistant gastric cancer cells, can be transferred to recipient cells sensitive to cisplatin via exosomes. The specific induction of gastric cancer cell apoptosis and cisplatin resistance indicates that inhibiting miR-769-5p may represent a potential therapeutic intervention strategy for the treatment of refractory gastric cancer.

## Materials and methods

All the materials and methods, and abbreviations are included in **Supplementary Materials and Methods**.

## Supplementary Materials and Methods

### Patient tissue and blood samples

Samples for cancer patients, including tissue and plasma specimens, were collected from the First Affiliated Hospital of Nanjing Medical University. Blood samples (serum) from 19 cisplatin-resistant patients and 41 cisplatin-sensitive patients were collected and stored at −80 °C. Other samples of 150 cases (75 pairs of GC tumor and normal tissuess) were embedded with 75 paraffin and analyzed by tissue microarray. Clinicopathological features, including age, sex, tumor site, tumor size, differentiation grade, Lauren classification, TNM stage (American Joint Committee on Cancer classification, AJCC), and lymphatic invasion, were also collected and analyzed (Table 1). This study was approved by the Ethics Committee of the First Affiliated Hospital of Nanjing Medical University. All patients signed an informed consent.

**Table 1.** Correlation of relative miR-769-5p expression with the clinicopathological characteristics of 150 patients with gastric cancer.

| Relationship between miR-769-5p expression and clinicopathologic factors of patients with gastric cancer |  |  |  |  |
| --- | --- | --- | --- | --- |
| Parameter | No. of patients | miR-769-5p(low) | miR-769-5p(high) | Pvalue (*P<0.05) |
| Sex |  |  |  |  |
| Male | 51 | 23 | 23 | 0.285 |
| Female | 24 | 14 | 14 |  |
| Age (year) |  |  |  |  |
| <60 | 60 | 33 | 27 | 0.133 |
| ≥60 | 15 | 5 | 10 |  |
| Tumor size (cm) |  |  |  |  |
| < 5 | 23 | 16 | 7 | 0.029 |
| ≥5 | 52 | 22 | 30 |  |
| Differentiation grade |  |  |  |  |
| well-moderate | 43 | 26 | 17 | 0.049 |
| poor-undifferentiation | 32 | 12 | 20 |  |
| T stage |  |  |  |  |
| T1-T2 | 7 | 6 | 1 | 0.051 |
| T3-T4 | 68 | 32 | 36 |  |
| Lymph node status |  |  |  |  |
| Negative | 23 | 17 | 6 | 0.007 |
| Positive | 52 | 21 | 31 |  |
| Distant metastasis |  |  |  |  |
| M0 | 75 | 38 | 37 |  |
| M1 | 0 | 0 | 0 |  |
| TNM stage |  |  |  |  |
| I-II | 31 | 21 | 10 | 0.013 |
| III-IV | 44 | 17 | 27 |  |

### Cell culture and treatment

The HEK-293T cell line was purchased from Type Culture Collection of the Chinese Academy of Sciences (Shanghai, China). Gastric cancer BGC823, SGC7901 cell lines, cisplatin-resistant BGC823/ DDP, and SGC7901/DDP cells were a kind gift from Professor Jianwei Zhou (School of Public Health, Nanjing Medical University). All cell lines were cultured in RPMI 1640 media (Gibco, Carlsbad, CA, USA) containing 10% fetal bovine serum (FBS) (ScienCell, CA, USA) and supplemented with 100 μg/ml streptomycin, 100 U/ml penicillin in a humidified atmosphere containing 5% CO2 at 37 °C. BGC823/ DDP and SGC7901/ DDP cells were cultured in a medium maintained with 0.5 μg/ml cisplatin (First Affiliated Hospital of Nanjing Medical University). Before the experiments, cell were cultured in a drug-free medium for at least 7 days. Cycloheximide (CHX)(Sigma-Aldrich, MO, USA) and MG132 (Selleck Chemicals, USA) were used at the indicated concentrations.

### Exosome isolation and characterization

Cell culture supernatant was collected after being washed with PBS and incubated with freshly prepared complete medium containing exosome-free FBS for 48h. Exosomes were isolated from the conditioned medium by differential centrifugation. Conditioned medium was centrifuged at 300 g for 10 min and then at 2,000 g for 20 min at 4 °C. The supernatant was then passed through a 0.22-μm filter (Millipore, Burlington, MA, USA) to remove shedding vesicles and other vesicles larger in size. Finally, the supernatant was centrifuged at 110,000 × g for 70 min. Pelleted exosomes were resuspended in PBS and collected by ultracentrifugation again at 100,000 g for 90 min (all steps were performed at 4 °C). Exosomes were collected from the pellet and resuspended in 100 μ L of PBS and subjected to several experiments. The fractionation and purification of exosomes from conditioned media (CMs) and blood serum were collected by ultracentrifugation (Beckman Coulter) and ExoQuick Exosome Precipitation Solution (SBI, CA, USA) respectively. Exosomes were then identified by Transmission Electron Microscope (TEM) (Philips TECNAI 20, Netherland), and their particle morphology and size were analyzed. The concentration and number of exosomes were detected by nanoparticle tracking analysis (NTA). Exosome protein markers were identified by Western blot assay and flow cytometry analysis (FACS Calibur, BD Biosciences, USA) .

### PKH26 Staining for Exosomes

The isolated exosomes were labeled with PKH26 Red Fluorescent Cell Linker Kits (Sigma). Exosomes were first resuspended in 100 μL Diluent C. A dye solution (4 × 10−6 M) was prepared by adding 0.4 μL PKH26 ethanolic dye solution to 100 μL Diluent C. The 100 μL exosome suspension was then mixed with the 100 μL dye solution by pipetting. After incubating the cell and dye suspension for 5 min with periodic mixing, the staining was stopped by adding 200 μL serum and incubating for 1 min. The stained exosomes were finally washed twice with 1× PBS, and they were resuspended in a fresh sterile conical polypropylene tube.

### Lentiviral, plasmid, and microRNA mimics/inhibitors package and cell transfection

The lentivirus encoding miR-769-5p overexpression or knockdown and negative control (769, NC, anti-769, anti-NC) were designed and produced by GENECHEM (Shanghai, China). The lentivirus were added to BGC823 BGC823/DDP, SGC7901 and SGC7901/DDP cells respectively and stable cell lines were obtained by selection with puromycin (Sigma-Aldrich, MO, USA). The infection efficiency was confirmed by fluorescence microscopy and real-time quantitative RT-PCR (qRT-PCR). pcDNA3.1 vector containing CASP9-wt, CASP9-mut, RNF20-wt or RNF20-mut, and a control vector were purchased from GENECHEM (Shanghai, China). miR-769-5p mimics, inhibitor and control, Cy3-miR-769-5p mimics and control were produced by GenePharma (Shanghai, China). Plasmids and miRNA mimics or inhibitors were transfected into cells with Lipofectamine 3000 (Invitrogen) according to the manufacturer’s instructions. The siRNAs and controls were designed and synthesized by RiboBio (Guangzhou, China). The siRNAs were transfected into the cells by DharmaFECT4 (Dharmacon, IL, USA); all sequences are listed in **Additional file 2 Table S1**.

### RNA extraction and quantitative RT-PCR

Total cellular and exosomal RNA was extracted from exosomes, co-cultured cells or GC cells, and frozen xenograft tumor tissues using TRIzol reagent (Invitrogen, CA, USA). Isolated RNA was used for the reverse transcription reaction with HiScript Q RT SuperMix for qPCR (Vazyme, Jiangsu, China). Quantitative RT-PCR was carried out with SYBR Green PCR Master Mix (Vazyme) using an ABI Prism 7900 Sequence detection system (Applied Biosystems, Canada). The relative expression of miR-769 was normalized to U6 levels, and CASP9, RNF20, p53 mRNA expression were normalized to GAPDH by qPCR using Power SYBR Green (Takara, Dalian, China). Data were calculated by the2 (−ΔΔCT) method. The related primers are synthesized by Ribobio (Guangzhou, China) and listed in **Additional file 2: Table S2**.

### Dual-luciferase reporter assays

293T cells (3 × 104 cells per well) were seeded onto 24-well plates 1 day before transfection and were co-transfected by Lipofectamine™ 3000 (Invitrogen, USA) with luciferase reporter (200 ng per well) using pmiR-REPORT™ luciferase vectors (pmirGLO) containing wild-type or mutant 3’-UTR of CASP9 and RNF20 and miR-769-5p mimics or miR-769-5p mimic-NC to examine the miRNA binding ability. The cells were washed and lysed with the passive lysis buffer from the Dual-Luciferase Reporter Assay System (Promega Corp). About 24 h later, a Dual-Luciferase Reporter Assay kit (Promega, USA) was used to measure the luciferase and renilla activity of these samples according to the manufacturer’s instructions. Relative luciferase activity was first normalized with Renilla luciferase activity and then compared with those of the respective control. Wild-type and mutated CASP9 or RNF20 3′ UTRs were synthesized and inserted into the p-MIR-REPORT plasmid by Genechem, Shanghai, China.

### Colony formation assay

GC cells (500 cells/well in six-well) were performed to detect the proliferation capacity. After incubation at 37 °C, 5% CO2 for two weeks, the plates were washed with PBS, fixed with 4% paraformaldehyde, stained with 0.1% crystal violet, washed three times with water, and analyzed. The assay was repeated three times in duplicate, and the numbers of colony formation counted.

### Cell viability assay

Cells (1 × 104/well) were seeded in 96-well plates and treated with cisplatin from 0.2 to 6.4 μg/ml for 24 h. A CCK-8 assay was performed to detect cells viability using a Cell Counting Kit 8 (Dojindo, Japan) and a OD450 nm (Synergy4; BioTek, Winooski, VT, USA). Based on protocols of CCK-8 kits cells were seeded, cultured for 24 h, and further cultured in 100 μL medium with 10 μL CCK-8 reagent. Absorbance at 450 nm was determined using a Multiscan FC plate reader (Thermo Fisher).

### Cell Migration Assay

The migratory capacity of GCs was tested by using a Transwell Boyden Chamber (6.5 mm, Costar) with polycarbonate membranes (8-μm pore size) on the bottom of the upper compartment. A total of 2 × 104 cells was suspended in serum-free media. Meanwhile, the lower chambers were loaded with 0.5 mL RPMI1640 containing 5% FBS, and the plates containing Transwell inserts were incubated. After incubation at 37 °C, 5% CO2 for 12 h, the upper chamber was cleaned with a cotton swab, and the lower chamber was washed with PBS. The cells that penetrated through the membrane were fixed with 90% ethanol for 15 min at room temperature, stained with 0.1% crystal violet solution, washed three times with water, and imaged by Inversion Microscope (Zeiss, Germany). The assay was repeated three times in duplicate. We obtained images of migrated cells by using a photomicroscope, and we quantified cell migration by blind counting with five fields per chamber.

### Apoptosis assay

The flow cytometry analysis was performed by Annexin V-APC/PI Apoptosis Detection Kit (Vazyme, Jiangsu, China) according to the manufacturer’s instructions.

The cells were analyzed with a BD FACS Calibur flow cytometer using CellQuest Pro software (FACS Calibur, BD Biosciences, USA).

### TUNEL assay

GC cells were fixed with paraformaldehyde for 30 min on ice. Then, terminal deoxynucleotidyl transferase dUTP nick end labeling (TUNEL) kit was used according to the manufacturer ’ s instructions (TUNEL BrightGreen Apoptosis Detection Kit, Vazyme, Jiangsu, China) and DAPI (4′,6-diamidino-2-phenylindole) was used for nuclear staining. TUNEL-positive areas were quantified under an Olympus FSX100 microscope (Olympus, Tokyo, Japan).

### Fluorescence assay

4′,6-diamidino-2-phenylindole (DAPI) (Invitrogen, USA) was used for cell nuclear staining. Rhodamine-conjugated secondary antibody (Cell Signaling Technology, USA) for γ-H2AX (1:250, Abcam, ab81299) protein and DAPI for nuclear staining. The slides were visualized for immunofluorescence with a laser scanning microscope (Zeiss, Germany).

### Western blot, immunohistochemistry (IHC), and immunoprecipitation (IP) assay

Cell or tissue samples were lysed by RIPA buffer mixed with protease and phosphatase inhibitor cocktails. Serum proteins were extracted with Serum Protein Extraction Kit (Qcheng Bio, China). The proteins were then separated by 10% SDS-PAGE and transferred onto PVDF membranes. Western blot assays were performed according to previously reported data [1]

The immune-complexes were detected with ECL Western Blotting Substrate (Thermo Fisher) and visualized with BIO-RAD (BIO-RAD Gel Doc XR+, USA). Immunohistochemistry and immunoprecipitation were done as previously reported [2]. Positive cells were counted in five random fields per slide. Primary antibodies and appropriate secondary antibodies used for the experiments are listed: TSG101 (1:1000, Abcam, ab125011), Calnexin (1:1000, Abcam, ab92573), CD81 (1:1000, Proteintech, 66866-1-Ig), CD63 (1:1000, Abcam, ab134045), γ -H2AX (1:250, Abcam, ab81299), caspase-9 (1:1000, CST, # 9504S), caspase-3 (1:1000, CST, # 9662), cleaved caspase-3 (1:1000, CST, # 9661), BAX (1:10000, Proteintech, 50599-2-Ig), Bcl-2 (1:1000, CST, #3498), p53 (1:5000, Proteintech, 10442-1-AP), EDD4L (1:5000, Proteintech, 13690-1-AP), RNF20(1:1000, Proteintech, 21625-1-AP), Ubiquitin(1:1000, CST, # 3936S), β-actin (1:1000, Beyotime, AF0003), GAPDH (1:1000, Beyotime, AF0006). Incubation with the goat anti-rabbit secondary antibody (1:1000, Beyotime, A0208) or the goat anti-mouse secondary antibody (1:1000, Beyotime, A0216).

### RNA in situ hybridization (ISH)

BaseScope™ Reagent Kit v2-RED (Advanced Cell Diagnostics, CA, USA) was used for ISH following the user manual. RNA in situ hybridization (ISH) was performed according to previously reported data . Standard RNAScope protocols were used according to manufacturer’s instructions and were performed according to previously reported data [3]. The following probes were used: miRNAscope Probe - SR-hsa-miR-769-5p-S1 (ACD; 1029501-S1), miRNAscope Positive Control Probe - SR-RNU6-S1 (ACD; 727871-S1), miRNAscope Negative Control Probe - SR-Scramble-S (ACD; 727881-S1).

### A nude mouse model

• 4- week-old (BALB/c) were obtained from Model Animal Research Center Of Nanjing University, China. All the animals were housed in an environment with a temperature of 22 ± 1 °C, relative humidity of 50 ± 1%, and a light/dark cycle of 12/12 hr and had access to water and food *at libitum*. All animal studies (including the mice euthanasia procedure) were done in compliance with the regulations and guidelines of Nanjing Medical University institutional animal care and conducted according to the AAALAC and the IACUC guidelines (IACUC-1902006)

a. Forty 4-week-old (BALB/c) male nude mice were randomly divided into two groups (20 mice in each group): BGC823+PBS and BGC823+BD EXO group. Briefly, 5 × 10^6^ BGC823 cells (100μL) were subcutaneously injected into the right flank of nude mice. When the average volume of nude mice reached approx. 50mm^3^, one group was intratumorally injected with BGC/DDP EXO (200ug/100μL cells per mouse) once every two days. When the tumor volume was 150-200mm^3^, each group were divided into two groups (10 mice in each group): BGC823+PBS+PBS, BGC823+PBS+DDP, BGC823+BD EXO+PBS and BGC823+BD EXO+DDP group, one group (BGC823+PBS+DDP, BGC823+BD EXO+DDP) was intraperitoneally injected with DDP (4mg/kg per mouse) every three days, and the other group (BGC823+PBS+PBS, BGC823+BD EXO+PBS) was injected with normal saline as the control group.
b. Forty 4-week-old (BALB/c) male nude mice were randomly divided into two groups (20 mice in each group): BGC NC and BGC 769. BGC823 cells with stable overexpression of miR-769-5p (BGC 769) and control cells (BGC NC) (5 × 10^6^/100μL cells per mouse) were subcutaneously injected into the right flank of nude mice. When the average volume of nude mice was about 150-200mm^3^, each group was divided into two groups: BGC NC+PBS, BGC 769+PBS, BGC NC+DDP and BGC 769+DDP. One group ( BGC/DDP anti769+DDP, BGC/DDP anti-NC+DDP) was intraperitoneally injected with DDP according to the standard of 4mg/kg every three days, and the other group (BGC NC+PBS, BGC 769+PBS) was intraperitoneally injected with normal saline as control.
c. Forty 4-week-old (BALB/c) male nude mice were randomly divided into two groups: BGC/DDPanti-769 and BGC/DDP anti-NC, with 20 mice in each group. BGC/DDP cells and control cells with stable knockdown expression of miR-769-5p (5 × 10^6^/100μL cells per mouse) were injected subcutaneously into the right flank of nude mice. When the average volume of nude mice was about 150-200mm^3^, each group was divided into two groups on average: BGC/DDPanti-769+PBS, BGC/DDP anti-NC+PBS, BGC/DDP anti769+DDP and BGC/DDP anti-NC+DDP. One group (BGC/DDP anti769+DDP, BGC/DDP anti-NC+DDP) was intraperitoneally injected with DDP according to the standard of 4mg/kg every three days, and the other group (BGC/DDPanti-769+PBS, BGC/DDP anti-NC+PBS) was intraperitoneally injected with normal saline as control.

Three weeks later, mice were sacrificed, and tumor tissues were prepared for histological examination: H&E staining, Western blot, and IHC assays. Tumor volume was measured using the following formula: *Tumor volume (mm^3^) = 0.5 ×width^2^ × length*.

### Statistical analysis

Statistical data were expressed as mean ± SD. One-way analysis of variance was used for three groups and more than three groups. All of the statistical analyses were assessed by software SPSS version 13.0 (SPSS, Chicago, IL, USA)and GraphPad Prism (GraphPad Software, Inc., SanDiego, CA, USA) software, comparisons among groups were done by the independent sample two-sided Student t-test. The ANOVA was performed to evaluate the statistical differences among groups. P- value of 0.05 or less was considered as statistical significance.

## Acknowledgements

We appreciate Prof. Jianwei Zhou for providing technical assistance

---

**Figure. S1.**
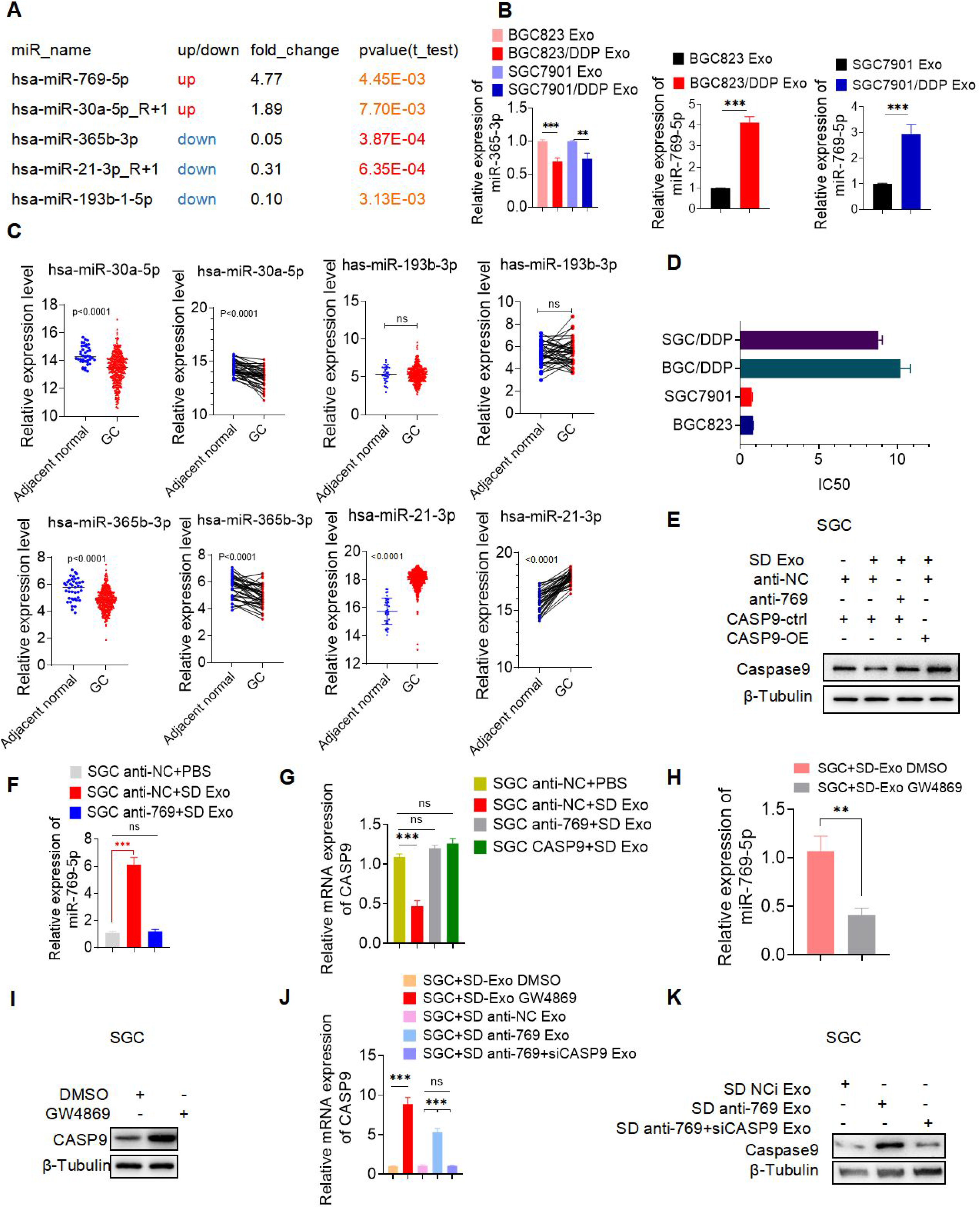
A. 5 most upregulated and downregulated miRNAs (hsa-miR-769-5p, hsa-miR-30a-5p, hsa-miR-365b-3p, hsa-miR-21-3p, hsa-miR-193b-5p) were selected based on the fold change and p value according to the result of differences in miRNAs expressed in two cell-derived exosome populations by using sequencing analysis. B. qPCR of miR-365-3p and miR-769-5p expression in exosomes released by BGC823, BGC823/DDP, SGC7901, SGC7901/DDP and found that the miR-769-5p expression was markedly higher in BD Exo and SD Exo. C. TCGA showed the expression of 4 miRNAs (hsa-miR-30a-5p, hsa-miR-365b-3p, hsa-miR-21-3p, hsa-miR-193b-5p) exculding miR-769-5p in GC and adjacent normal. D. IC50 of BGC823, BGC823/DDP, SGC7901, SGC7901/DDP cell lines. E. (relatead to Figure.2L) , G. (relatead to Figure.2K) The mRNA and protein levels of CASP9 in SGC anti-NC+ PBS, SGC anti-NC + SD Exo and SGC anti-769 + SD Exo. F. (relatead to Figure.2J) qRT-PCR showed the expression of miR-769-5p in in SGC anti-NC + PBS, SGC anti-NC + SD Exo and SGC anti-769 + SD Exo. H. (relatead to Figure.2N), I. (relatead to Figure.2M) RT-PCR and Western blot showed the expression of miR-769-5p in SGC + SD Exo DMSO and SGC + SD Exo GW4869. J. (relatead to Figure.2O), K. (relatead to Figure.2P) The upregulation of CASP9 mRNA and protein was detected by qRT-PCR and Western blot in SGC + SD Exo GW4869 compared to SGC + SD anti-769 Exo. Quantitative data from three independent experiments are shown as the mean ± SD (error bars). *P < 0.05, **P < 0.01, ***P < 0.001 (Student’s t-test)

**Figure. S2.**
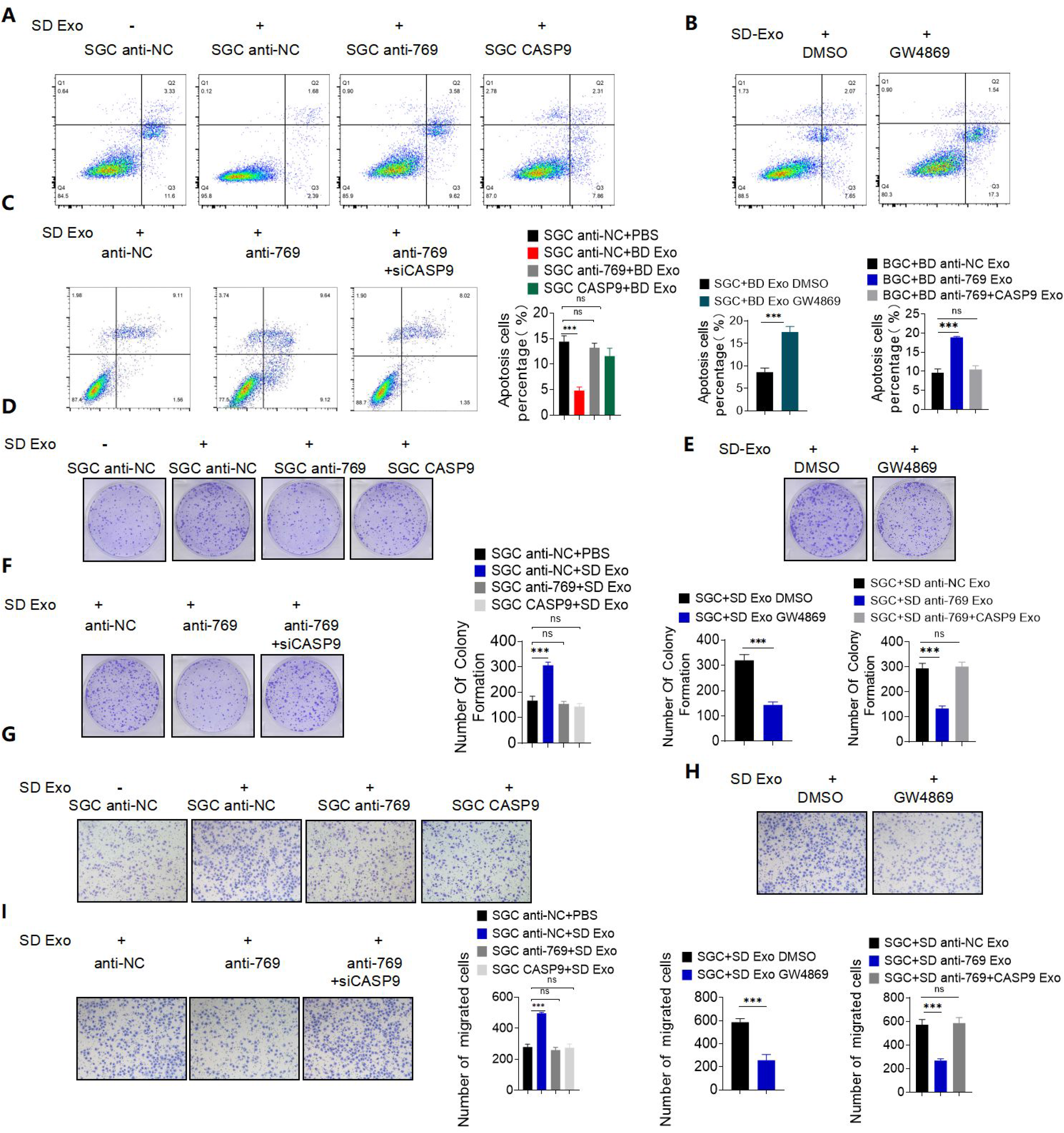
A. (relatead to Figure.3A) Flow cytometry assay detected cell apoptosis rate of SGC anti-NC + PBS, SGC anti-NC + SD Exo, SGC anti-769 + SD Exo and SGC CASP9 + SD Exo. B. (relatead to Figure.3B) Flow cytometry assay detected cell apoptosis rate of SGC + SD Exo DMSO and SGC + SD Exo GW4869. C. (relatead to Figure.3C) Flow cytometry assay detected cell apoptosis rate of SGC + SD anti-NC Exo, SGC + SD anti-769 Exo and SGC + SD anti-769 + siCASP9 Exo. D. (relatead to Figure.4C) The average colony numbers of three independent experiments were calculated in SGC anti-NC + PBS, SGC anti-NC + SD Exo, SGC anti-769 + SD Exo and SGCCASP9 + SD Exo. E. (relatead to Figure.4D) The average colony numbers of three independent experiments were calculated in BGC + SD Exo DMSO and BGC + SD Exo GW4869. F. (relatead to Figure.4E) The average colony numbers of three independent experiments were calculated in BGC + SD anti-NC Exo, BGC + SD anti-769 Exo and BGC + SD anti-769 + siCASP9 Exo. G. (relatead to Figure.4F) Migration and invasion ability of SGC anti-NC + PBS, SGC anti-NC+SD Exo, SGC anti-769+ SD Exo and SGCCASP9 + SD Exo were assessed by Transwell assay. H. (relatead to Figure.4G) Migration and invasion ability of BGC + SD Exo DMSO and BGC + SD Exo GW4869 were assessed by Transwell assay. I. (relatead to Figure.4H) Migration and invasion ability of BGC + SD anti-NC Exo, BGC + SD anti-769 Exo and BGC + SD anti-769 + siCASP9 Exo were assessed by Transwell assay. Quantitative data from three independent experiments are shown as the mean ± SD (error bars). *P < 0.05, **P < 0.01, ***P < 0.001 (Student’s t-test)

**Figure. S3.**
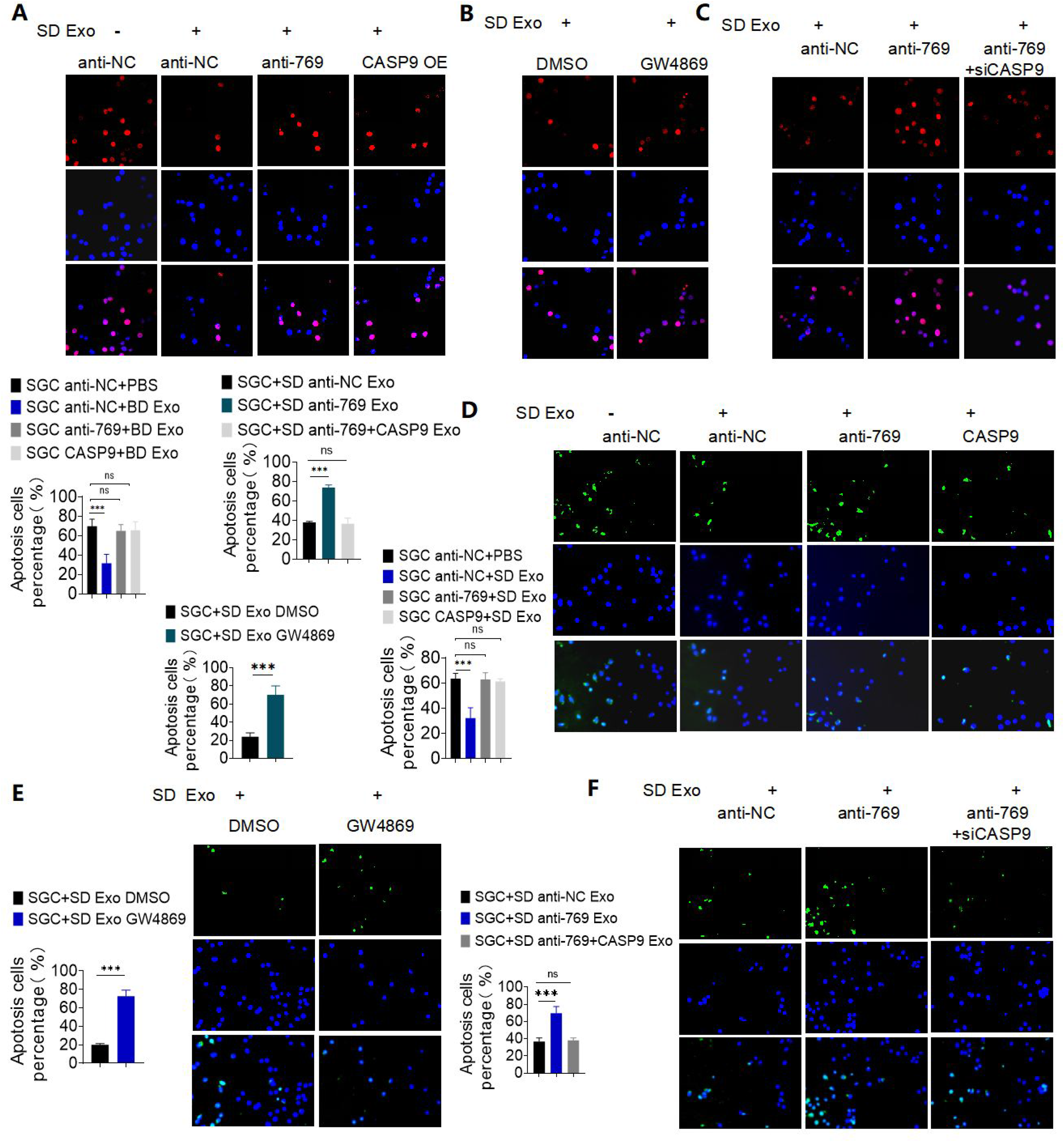
A. (relatead to Figure.3D) The level of γ -H2AX nuclear foci in SGC anti-NC + PBS, SGC anti-NC + SD Exo, SGC anti-769 + SD Exo and SGC CASP9+ SD Exo. B. (relatead to Figure.3E) The level of γ-H2AX nuclear foci in SGC + SD Exo DMSO and SGC + SD Exo GW4869. C. (relatead to Figure.3F) The level of γ-H2AX nuclear foci in SGC + SD anti-NC Exo, SGC + SD anti-769 Exo and SGC + SD anti-769 + siCASP9 Exo. D. (relatead to Figure.3G) TUNEL analysis detected cell apoptosis rate of SGC anti-NC + PBS, SGC anti-NC + SD Exo, SGC anti-769 + SD Exo and SGC CASP9 + SD Exo. E. (relatead to Figure.4A) TUNEL analysis detected cell apoptosis rate of BGC + SD Exo DMSO and BGC + SD Exo GW4869. G. (relatead to Figure.4B) TUNEL analysis detected cell apoptosis rate of BGC + SD anti-NC Exo, BGC + SD anti-769 Exo and BGC + SD anti-769 + siCASP9 Exo. Quantitative data from three independent experiments are shown as the mean ± SD (error bars). *P < 0.05, **P < 0.01, ***P < 0.001 (Student’s t-test)

**Figure. S4.**
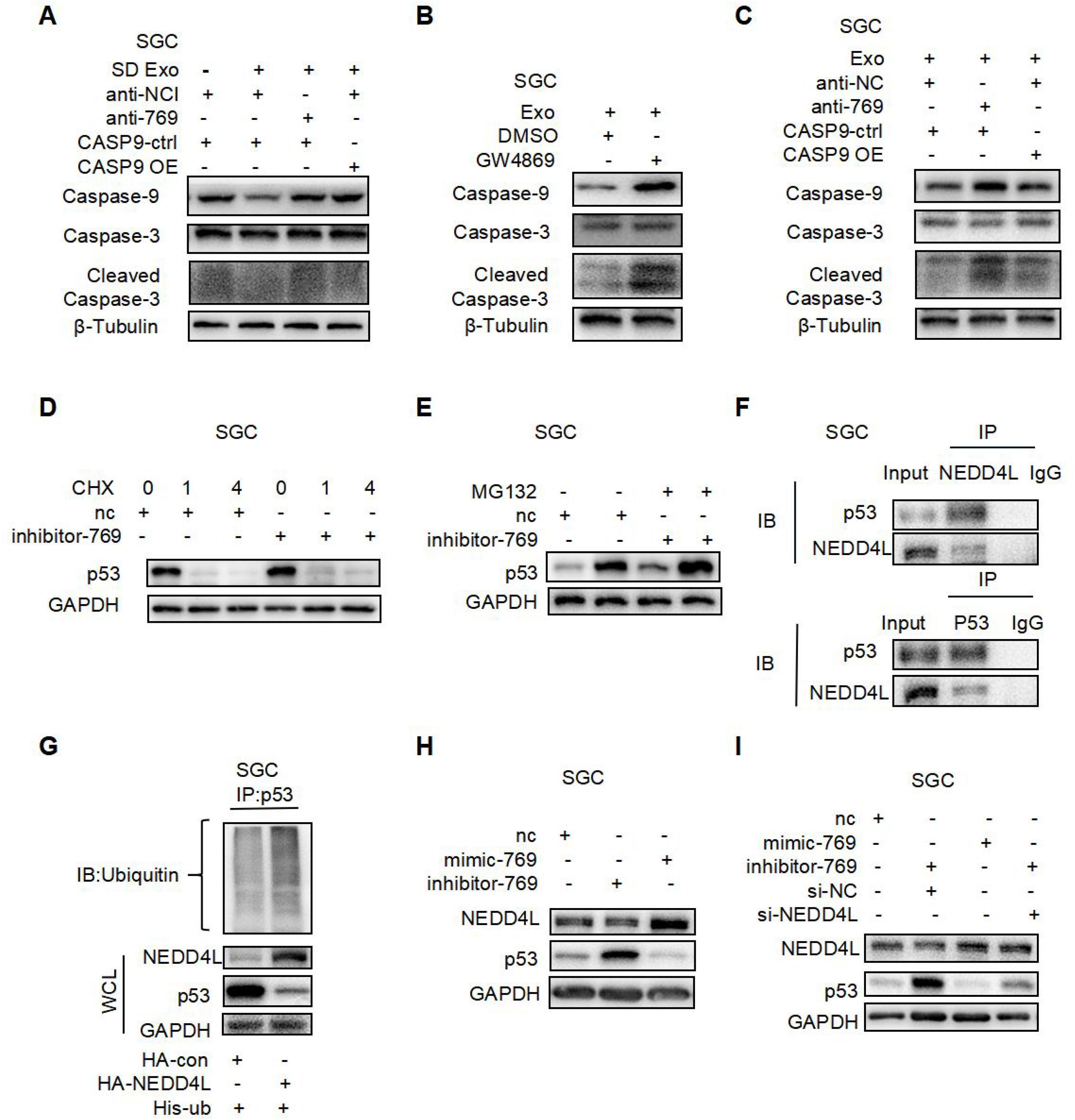
A. (relatead to Figure.5A) Western blot anaysis of caspase9, caspase3 and cleaved caspase3 in SGC + SD anti-NC Exo, SGC + SD anti-769 Exo and SGC + SD anti-769 + siCASP9 Exo. B. (relatead to Figure.5B) Western blot anaysis of caspase9, caspase3 and cleaved caspase3 in SGC + SD Exo DMSO and SGC + SD Exo GW4869. C. (relatead to Figure.5C) Western blot anysis of caspase9, caspase3 and cleaved caspase3 in SGC + SD anti-NC Exo, SGC + SD anti-769 Exo and SGC +SD anti-769+siCASP9 Exo. D. (relatead to Figure.6E) Western blot analysis of p53 protein level of 100ug/ml treated with cycloheximide (CHX) changes with treatment time. E. (relatead to Figure.6F)Western blot analysis of p53 protein level after MG-132 (10um) treatment. F. (relatead to Figure.6H) Co-IP detected the interaction between NEDD4L and p53 in SGC cells. I. Co-IP and western blot detected. G. (relatead to Figure.6I) p53 ubiquitination modification mediated by NEDD4L. H. (relatead to Figure.6J), I. (relatead to Figure.6K) The expression of NEDD4L and p53 protein levels when miR-769-5p is knocked down or overexpressed. Quantitative data from three independent experiments are shown as the mean ± SD (error bars). *P < 0.05, **P < 0.01, ***P < 0.001 (Student’s t-test)

**Figure. S5.**
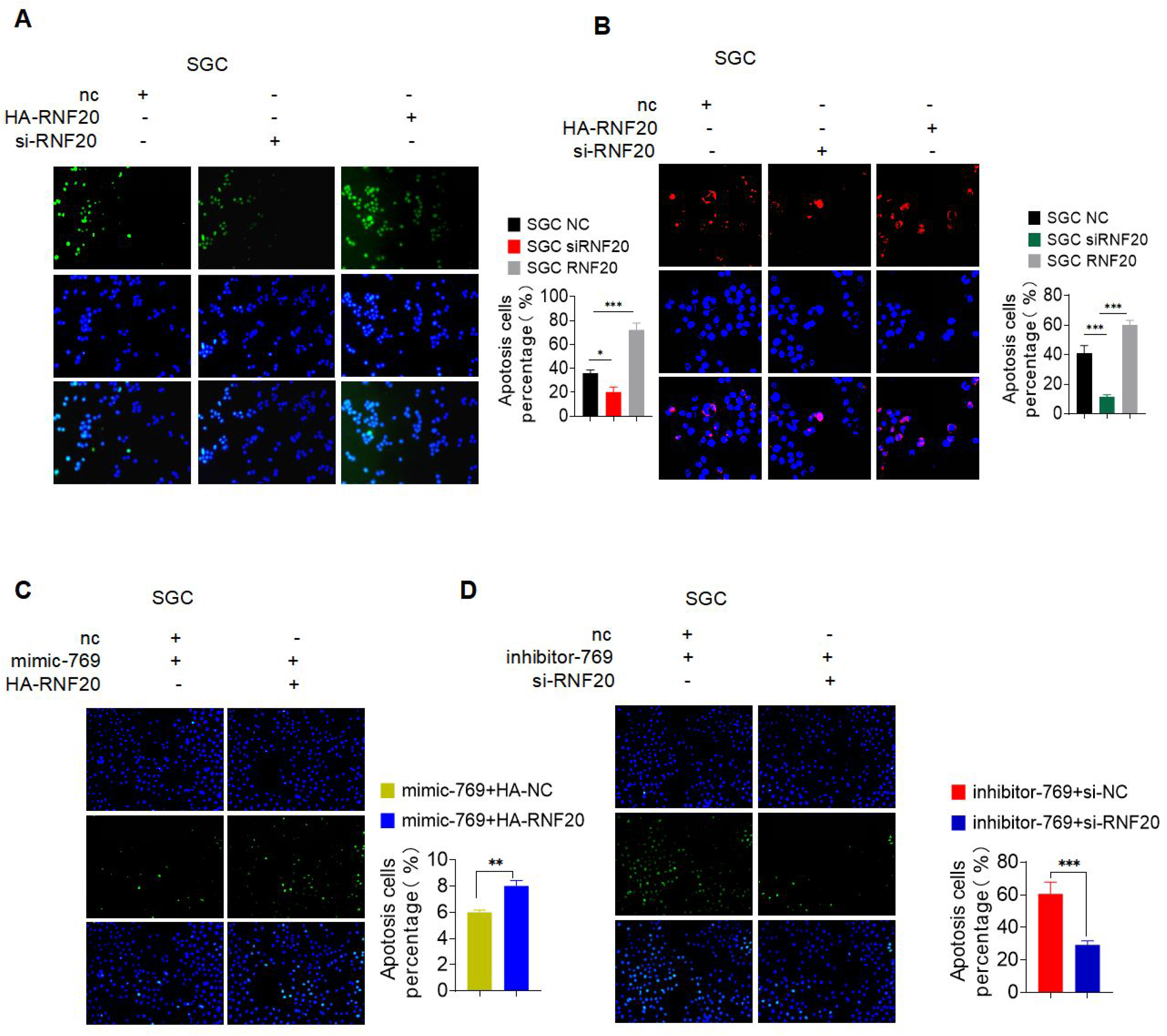
A. (relatead to Figure.7A) TUNEL analysis detected cell apoptosis rate of SGC NC, SGC HA-RNF20 and SGC si-RNF20. B. (relatead to Figure.7B) The level of γ -H2AX nuclear foci in SGC NC, SGC HA-RNF20 and SGC si-RNF20. C (relatead to Figure.7D), D (relatead to Figure.7E). The recovery proved that miR-769-5p inhibits the process of apoptosis by down-regulating RNF20 by analysis of TUNEL. Quantitative data from three independent experiments are shown as the mean ± SD (error bars). *P < 0.05, **P < 0.01, ***P < 0.001 (Student’s t-test)

**Figure. S6.**
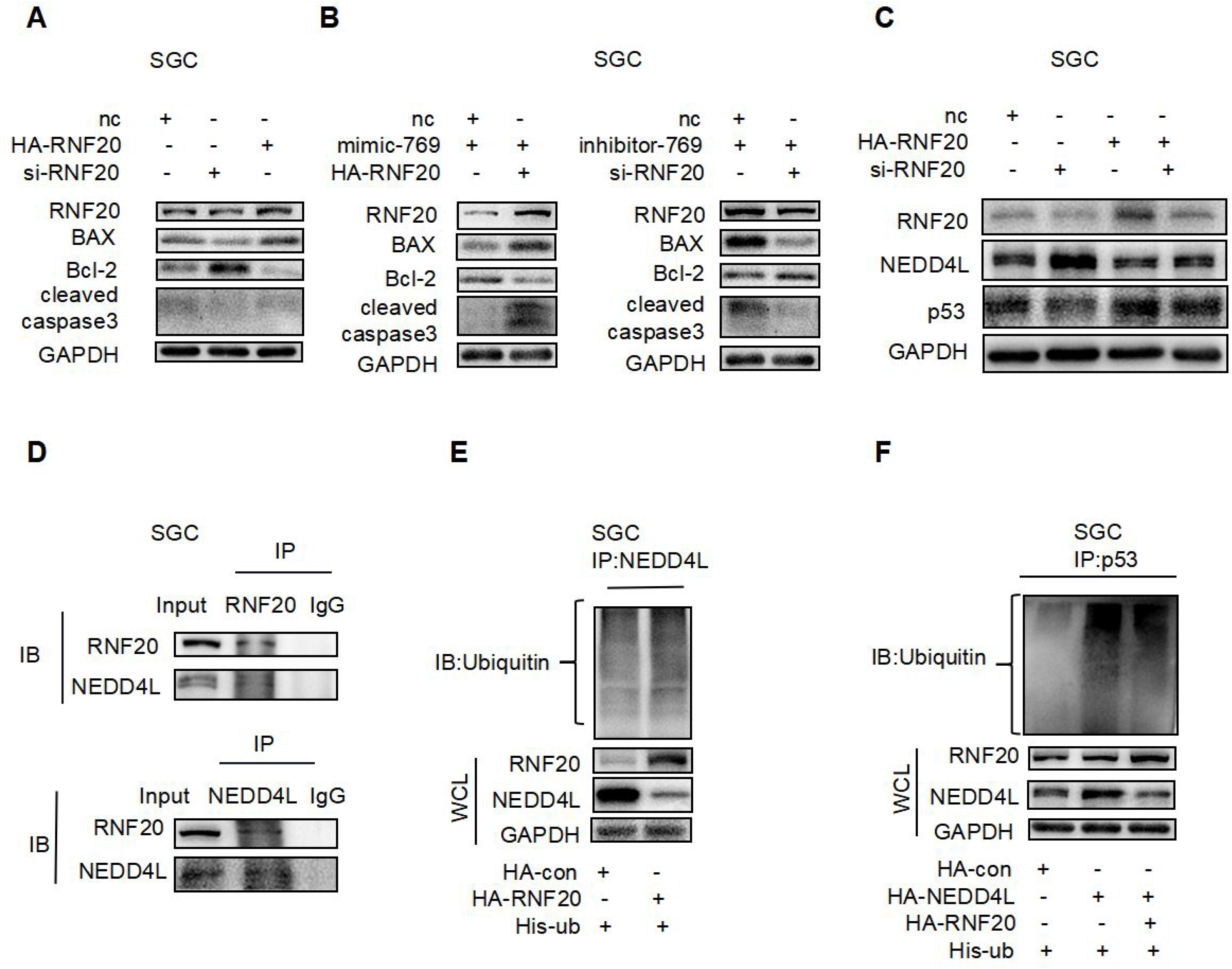
A. (relatead to Figure.7C), B (relatead to Figure.7F) The western blot analysis of Bax, Bcl-2 and cleaved caspase 3 proved the mediation of RNF20 on apoptosis. C. (relatead to Figure.7G) The protein levels of NEDD4L and p53 when RNF20 overexpression and knockdown. D. (relatead to Figure.7I) Co-IP proved that NEDD4L interacts with RNF20. E. (relatead to Figure.7J), F (relatead to Figure.7K) Co-IP proved that the ubiquitination modification of NEDD4L is mediated by RNF20. Quantitative data from three independent experiments are shown as the mean ± SD (error bars). *P < 0.05, **P < 0.01, ***P < 0.001 (Student’s t-test)

**Additional file 2: Table S1.**
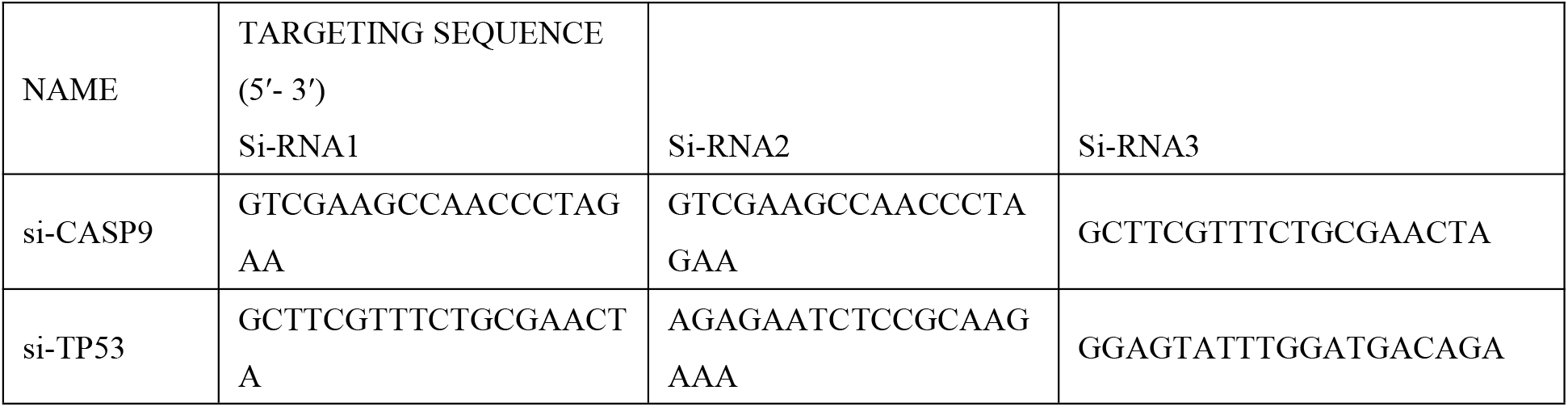
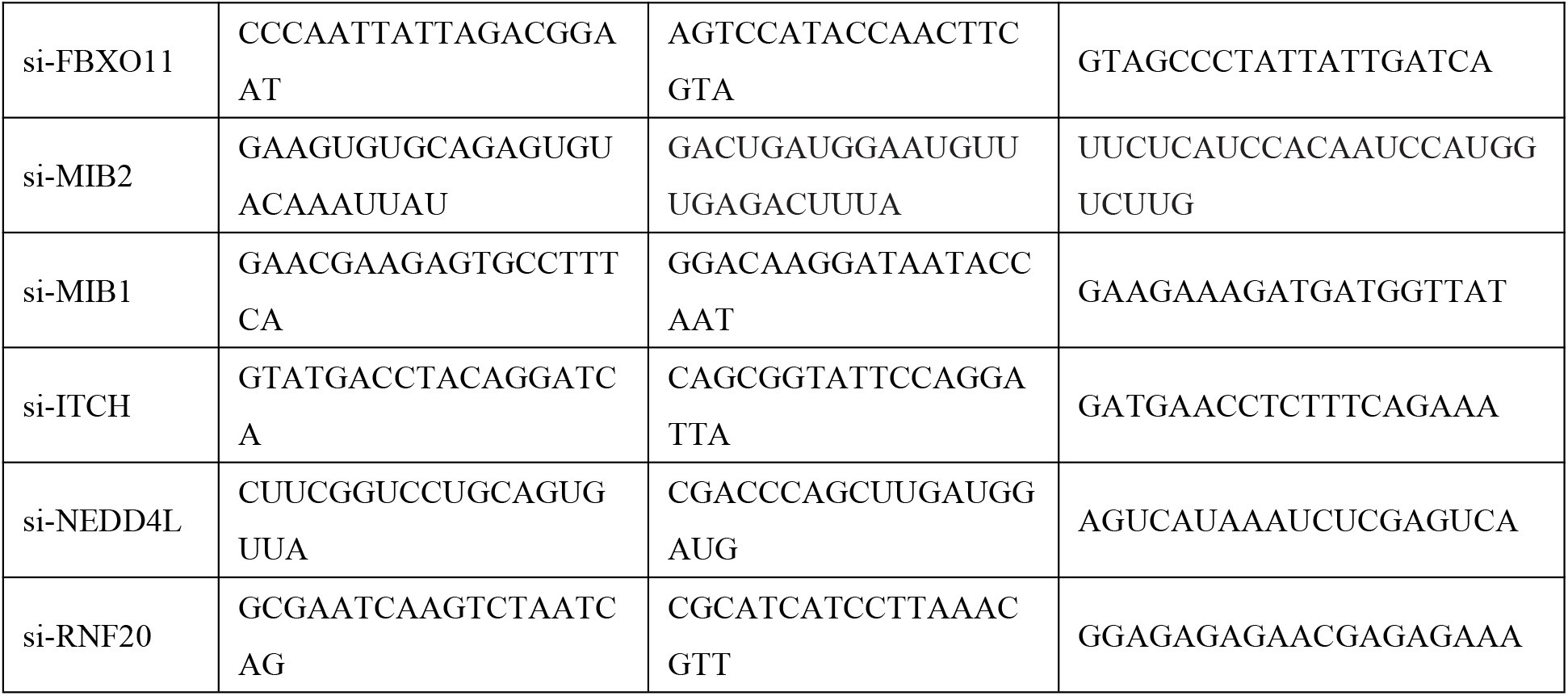
All sequences of siRNAs are listed.

**Additional file 2: Table S2.**
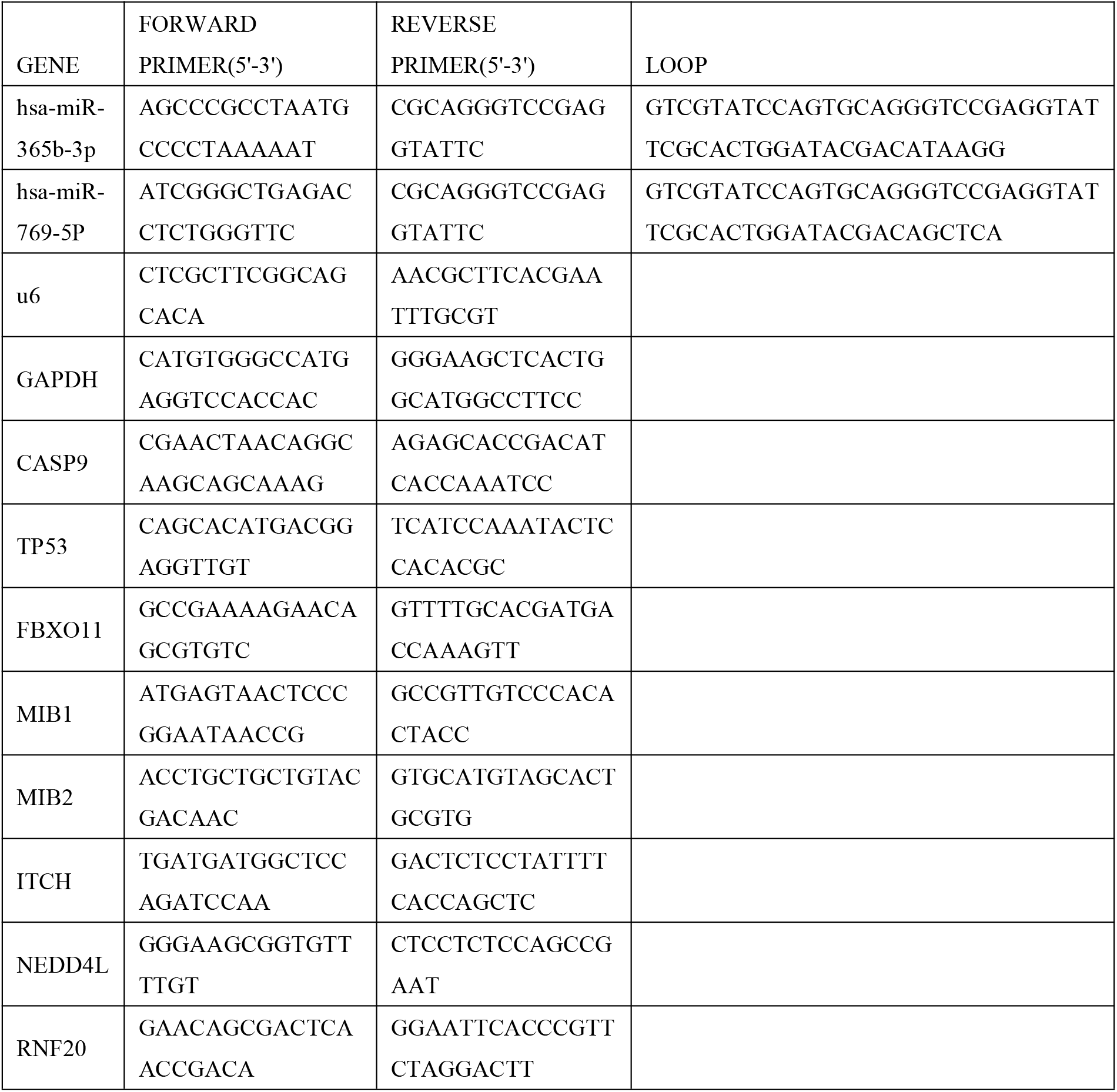
The related primers are synthesized.

## Abbreviations

GC: gastric cancer
miRNA: MicroRNA
UPS: ubiquitin-proteasome system
NTA: NanoSight particle tracking analysis
FCM: flow cytometry
BD Exo: BGC823/DDP secreted exosomes
BC Exo: BGC secreted exosomes
TME: tumor microenvironment
BGC anti-769: BGC823 cells with lentiviral vectors stably expressing miR-769-5p inhibitor
BGC anti-NC: negative control miRNA inhibitor
BGC CASP9: BGC823 cells with lentiviral vectors stably expressing CASP9
CHX: cycloheximide
siRNA: small interfering RNA
CoIP: Co-immunoprecipitation

